# A multidisciplinary analysis of the as-isolated *Escherichia coli* SufBC_2_D complex reveals the presence of two new iron-sulfur clusters

**DOI:** 10.1101/2022.03.29.486248

**Authors:** Giulia Veronesi, Julien Pérard, Martin Clémancey, Catherine Gerez, Yohann Duverger, Isabelle Kieffer, Frédéric Barras, Serge Gambarelli, Geneviève Blondin, Sandrine Ollagnier de Choudens

## Abstract

Iron-sulfur (Fe-S) clusters are essential inorganic cofactors dedicated to a wide range of biological functions including electron transfer and catalysis. Specialized multi-protein machineries present in all types of organisms support their biosynthesis. These machineries encompass a scaffold protein on which Fe-S clusters are assembled before being transferred to cellular targets. Here, we describe the first characterization of the native Fe-S cluster of the anaerobically purified SufBC_2_D scaffold from *Escherichia coli* by XAS, Mössbauer, UV-visible absorption and EPR spectroscopy. Interestingly, we propose that SufBC_2_D harbors two types of Fe-S cluster, a [2Fe-2S] cluster with an unprecedented usual coordination and a previously unreported [3Fe-3S] cluster. These data combined with mutagenesis and biochemistry allow to propose ligands for these clusters. These results support the hypothesis that both SufB and SufD are involved in Fe-S cluster ligation and are discussed in the context of Fe-S cluster biogenesis where both [2Fe-2S] and [4Fe-4S] clusters need to mature cellular Fe-S protein targets.

## INTRODUCTION

Iron-sulfur clusters are essential protein cofactors that enable proteins to perform a variety of unique and complex functions, ranging from redox chemistry, amino-acid biosynthesis to DNA replication and repair.^1–3^ These cofactors exist under different forms, [2Fe-2S], [3Fe-4S] and [4Fe-4S] being the most prevalent ones. Their biogenesis is highly regulated and conserved among organisms. Despite their significance, a number of questions still remains regarding the mechanism through which Fe-S clusters are generated. In *Escherichia coli* (*E. coli*), production of Fe-S clusters occurs via two major pathways, Isc and Suf.^4^ While Isc directs Fe-S generation under normal conditions, Suf takes over during periods of iron depletion and oxidative stress. The *E. coli* Suf system consists of six proteins (SufABCDSE) among which SufB, SufC and SufD form the SufBC_2_D complex, which serves as a scaffold for the synthesis of nascent Fe-S clusters.^5^ The cysteine desulfurase SufS uses its PLP to mobilize sulfur from L-cysteine; SufE enhances SufS cysteine desulfurase activity^6, 7^ while SufA, an Fe-S carrier, transports Fe-S clusters from SufBC_2_D to targets.^8, 9^

The crystal structure of *E. coli* SufBC_2_D without Fe-S cluster was solved at 2.9 angstrom.^10^ SufB interacts with both SufD and SufC, and SufD with both SufB and SufC. Thus, each SufC subunit is bound to a subunit of SufB-SufD heterodimer. After chemical reconstitution or anaerobic purification of SufBC_2_D, the complex binds either a [2Fe-2S] or a [4Fe-4S] cluster.^5, 10–12^ It has been suggested for a long time that the cluster is localized on SufB subunit since *in vitro* SufB protein assembles after reconstitution an Fe-S cluster (either [2Fe-2S] or [4Fe-4S]) resembling that of the SufBC_2_D complex.^11, 13, 14^ However, recently *in vivo* data (color of host cells expressing wt vs variants SufBC_2_D) identified amino-acids both in SufB and SufD subunits which appeared to be important for the formation and/or coordination of the Fe-S cluster within the complex^15^, namely SufB^C405^, SufB^E432^, SufB^H433^, SufB^E434^, SufD^H360^ and SufD^C358^, supporting the notion that the Fe-S cluster is bound at the SufD-SufB interface. In order to solve this conundrum and gain insight in the actual organization of the Fe-S bound SufBC_2_D complex, we undertook a thorough characterization of the as-isolated *E. coli* Fe-S-SufBC_2_D complex by using X-ray absorption, Mössbauer, UV-visible and EPR spectroscopies, combined with biochemical and mutagenesis studies. Such a broad and multidisciplinary approach led us to propose two populations of Fe-S bound SufBC_2_D complexes, both tentatively assigned to new types of clusters.

## RESULTS and DISCUSSION

### XAS analysis of the anaerobically purified SufBC_2_D complex

His_6_-SufBC_2_D complex was co-synthesized with SufSE from the pETDuet-1 plasmid.^11^ After anaerobic purification, SEC-MALS-RI confirmed the homogeneity of the sample with a BC_2_D stoichiometry (Figure S1). The as-purified SufBC_2_D complex contains an average of 2.4±0.2 Fe and 2.3±0.1 S per complex and displays a brown-blackish color with an unusual UV-vis absorption spectrum containing a main absorption band at 420 nm and shoulders at 320 nm, 530 and 620 nm (Figure 1B). UV-vis. spectrum is suggestive of a linear [3Fe-4S] cluster with absorptions at 320, 415, 513 and 573 nm.^16–18^ This complex was therefore analyzed by X-ray absorption spectroscopy. The X-ray Absorption Near Edge Structure (XANES) spectrum of the complex (Figure 2A) exhibits all the features associated to tetrahedral Fe in Fe-S clusters, notably a prominent pre-peak at ~7113 eV, a shoulder in its rising edge at ~ 7122 eV, and a relatively featureless post-edge trend. However, the XANES fingerprint alone does not allow us to discriminate between [4Fe-4S], [3Fe-4S] and [2Fe-2S], considering that such features have been observed for all kinds of clusters.^19–25^

**Figure 1.**
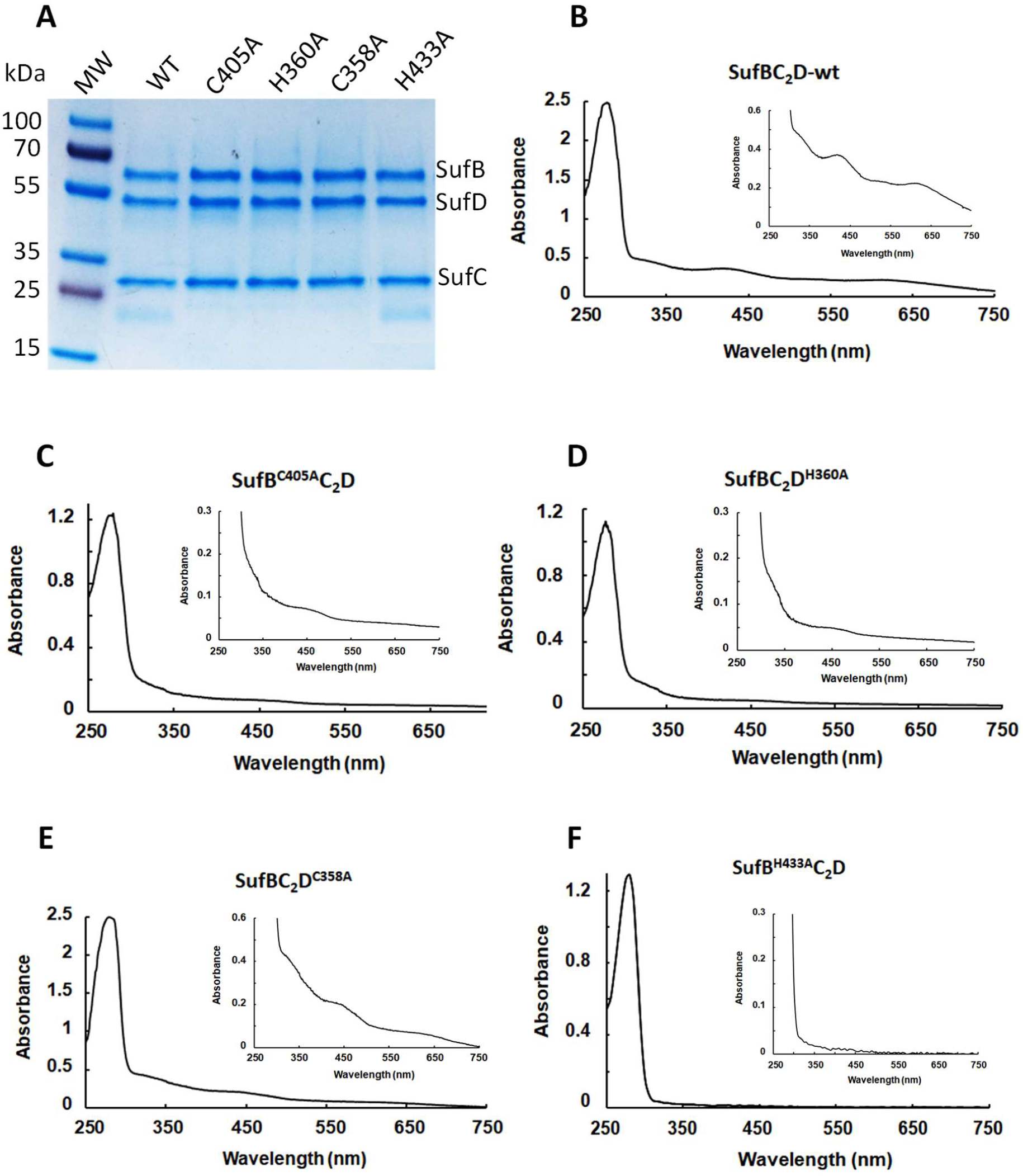
Anaerobic purification of SufBC_2_D-His wild-type and variants. SDS-PAGE (A) and UV-Vis absorption spectra ((B) to (F)) of anaerobically purified SufBC_2_D-WT and SufBC_2_D variants. The *insets* of UV-vis. spectra show an enhancement of the Fe-S absorption bands. MW: Molecular weight (kDa).

**Figure 2.**
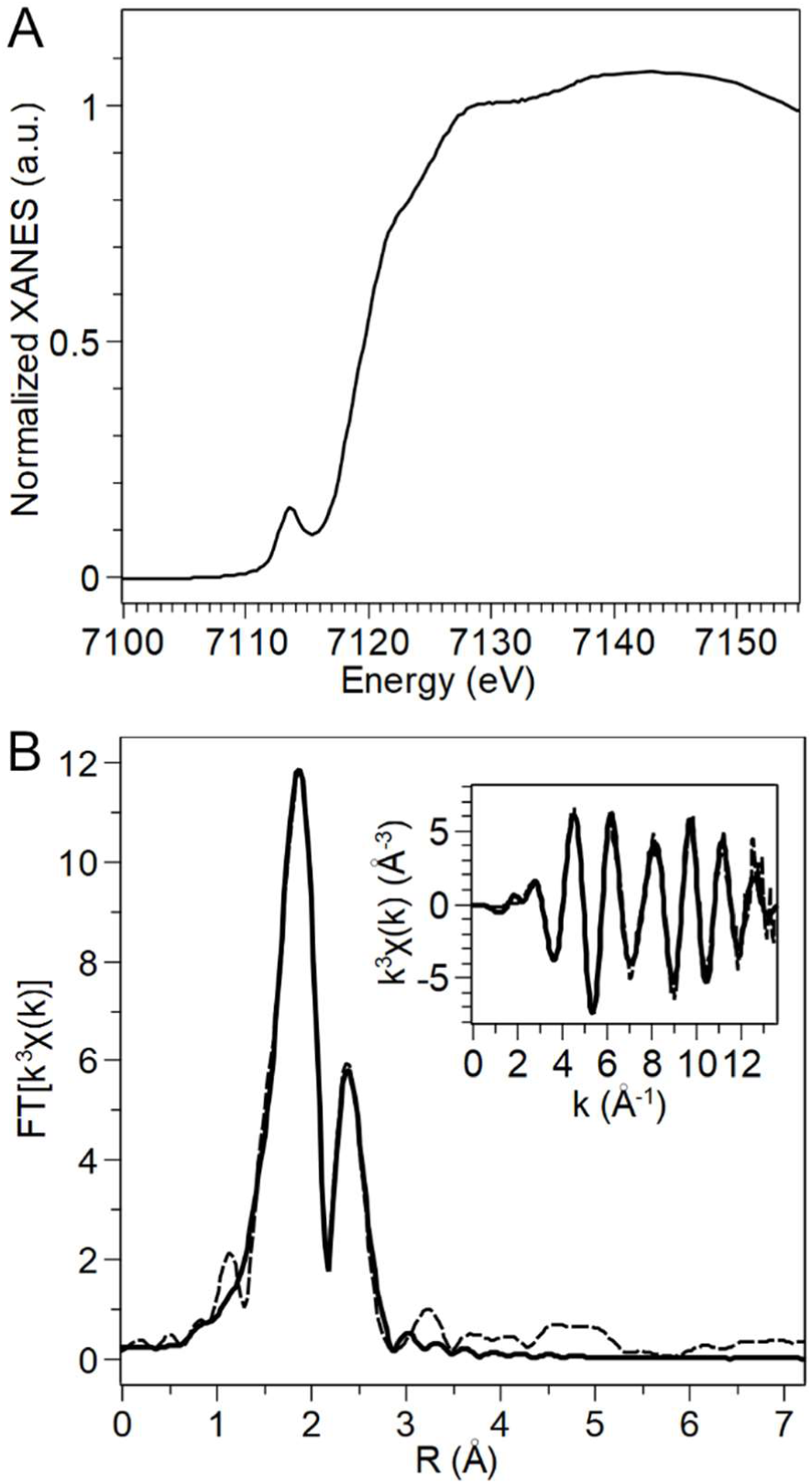
Fe K-edge X-ray Absorption Spectroscopy of the SufBC_2_D complex. (A) Experimental XANES spectrum. (B) Experimental (dashed curves) EXAFS spectrum in the reciprocal space (inset) and Fourier-transformed into the real space (main panel); relative best-fitting curve (solid curves) obtained in the real space (main panel), and back-transformed into the reciprocal space (inset).

In order to disclose the nature of the cluster, the Extended X-ray Absorption Near Edge Structure (EXAFS) spectrum was fitted in the range 1-3.0 Å, assuming a tetrahedral Fe environment composed of sulfur and oxygen/nitrogen ligands, and a variable number of second-shell Fe atoms (Figure 2B). In the first instance the number of ligands of each species, the relative distance, and Debye-Waller factors (σ^2^) were allowed to vary. The best-fitting average Fe environment includes 2.8 ± 0.3 S at 2.27 ± 0.01 Å from the absorber, 1.2 ± 0.3 N/O at 2.04 ± 0.02 Å, and 1.3 ± 0.3 Fe at 2.75 ± 0.01 Å. The σ^2^ values were 4.6 ± 0.9 ·10^−3^Å^2^, 9 ± 5 ·10^−3^Å^2^, and 5 ± 2 ·10^−3^Å^2^, respectively, and the goodness-of-fit R=0.6% (Figure 2B). The number of Fe around the absorber, 1.3 ± 0.3 Fe, is consistent with a [2Fe-2S] cluster or a mixture of clusters of higher nuclearity. However, the well-known correlation between the number of atoms of a given species around the absorber and the relative Debye-Waller factor could bias the fit results. Therefore, in order to disentangle these parameters, we built different models with fixed number of neighbors according to the existing Fe-S clusters, and fitted them to the experimental data by allowing only interatomic distances and Debye-Waller factors to vary. The models included *n* sulfur ligands and (4-*n*) oxygen/nitrogen ligands in the first shell around the absorber, and different numbers of Fe atoms in the second shell. The number of S atoms per Fe absorber, *n*, ranged from 2 (the minimum number of S ligands, encountered in a [2Fe-2S] cluster anchored to the protein through non-Cys residues) to 4 (in case the cluster is anchored to the protein through Cys residues only). Non-integer *n* values account for a possible asymmetry between different Fe centers in the cluster. The number of second-shell Fe atoms was fixed to 1, 1.33, 2, 3 to model [2Fe-2S], linear [3Fe-4S], cubane [3Fe-4S], and [4Fe-4S] clusters, respectively. This resulted in nineteen models to fit to the experimental data, reported in Table 1. The possible clusters corresponding to each model are represented in Figure S2. The comparison between the goodness-of-fit (R_fit_ values in Table 1) of [2Fe-2S] models, [3Fe-4S] models and [4Fe-4S] models with the same Fe coordination sphere (e.g. model 4S 1Fe vs 4S 3Fe) shows that the [2Fe-2S] and linear [3Fe-4S] clusters are systematically favored. Moreover, in [4Fe-4S] models, the dynamical parameter σ^2^ relative to the Fe-Fe interaction is always estimated as 0.013 Å^2^, a very high value for a metal-metal interaction, clearly overestimated to compensate for the too high value of Fe-Fe interactions in the model. [2Fe-2S] models and linear [3Fe-4S], instead, provide σ^2^ values between 0.003 and 0.005 Å^2^ for the same interaction, compatible with the values provided in literature for various Fe-S cluster systems.^20, 23, 25–27^

**Table 1.** Results of Ab initio Fitting of the Fe K-Edge EXAFS spectrum of the SufBC2D complex based on different cluster models.^a,b^

| Model | number<br>of Cys<br>ligands | Fe-S | | Fe-N/O | | Fe-Fe | | $\Delta E_0$<br>(eV) | $R_{\text{fit}}$<br>(%) |
| --- | --- | --- | --- | --- | --- | --- | --- | --- | --- |
| | | R<br>(Å) | $\sigma^2$<br>(10<sup>3</sup> Å<sup>2</sup>) | R<br>(Å) | $\sigma^2$<br>(10<sup>3</sup> Å<sup>2</sup>) | R<br>(Å) | $\sigma^2$<br>(10<sup>3</sup> Å<sup>2</sup>) | | |
| [2Fe 2S] models |  |  |  |  |  |  |  |  |  |
| 4S 1Fe | 4 | 2.263 (12) | 7.3 (9) | - | - | 2.731 (18) | 3 (2) | -3 (1) | 2.9 |
| 3.5S 0.5O/N<br>1Fe | 3 | 2.261 (12) | 6.0 (9) | 2.11 (7) | 5<sup>c</sup> | 2.733 (12) | 3.2 (11) | -2 (1) | 1.0 |
| 3S 1O/N 1Fe | 2 | 2.271 (7) | 5.1 (6) | 2.05 (2) | 8 (5) | 2.744 (9) | 3.2 (9) | -0.5<br>(8) | 0.6 |
| 2.5S 1.5O/N<br>1Fe | 1 | 2.281 (7) | 3.9 (6) | 2.03 (3) | 9 (3) | 2.753 (10) | 3.3 (10) | 1.2 (9) | 0.9 |
| 2S 2O/N 1Fe | 0 | 2.289 (11) | 2.8 (9) | 2.03 (2) | 8 (3) | 2.762 (15) | 3.6 (16) | 3 (1) | 1.8 |
| linear [3Fe 4S] models |  |  |  |  |  |  |  |  |  |
| 4S 1.33Fe | 4 | 2.262 (13) | 7.1 (9) | - | - | 2.736 (18) | 5 (2) | -3 (1) | 3.4 |
| 3.67S<br>0.33O/N<br>1.33 Fe | 3 | 2.260 (11) | 6.1 (20) | 2.16<br>(30) | 5<sup>c</sup> | 2.733 (20) | 5.2 (16) | -3 (2) | 1.5 |
| 3.33S<br>0.67O/N<br>1.33 Fe | 2 | 2.262 (8) | 5.6 (11) | 2.09 (7) | 8 (14) | 2.738 (13) | 5.2 (11) | -2 (1) | 0.9 |
| 3S<br>1O/N<br>1.33Fe | 1 | 2.269 (6) | 5.0 (6) | 2.05<br>(12) | 9 (6) | 2.745 (9) | 5.1 (9) | -0.8<br>(8) | 0.6 |
| 2.67S<br>1.33O/N<br>1.33Fe | 0 | 2.275 (6) | 4.2 (5) | 2.03 (1) | 10 (4) | 2.751 (9) | 5.2 (9) | 0.3 (7) | 0.6 |
| cubane [3Fe 4S] models |  |  |  |  |  |  |  |  |  |
| 4S 2Fe | 3 | 2.261 (15) | 6.8 (9) | - | - | 2.74 (2) | 8 (3) | -3 (1) | 4.9 |
| 3.67S<br>0.33O/N<br>2Fe | 2 | 2.259 (11) | 6 (3) | 2.2 (5) | 5<sup>c</sup> | 2.74 (3) | 9 (2) | -3 (2) | 2.5 |
| 3.33S<br>0.67O/N<br>2Fe | 1 | 2.259 (11) | 5.5 (8) | 2.11 (6) | 5<sup>c</sup> | 2.741 (14) | 8.7 (15) | -2 (1) | 1.8 |
| 3S 1O/N<br>2Fe | 0 | 2.265 (8) | 4.7 (7) | 2.06 (3) | 12 (10) | 2.745 (12) | 8.7 (12) | -1 (1) | 1.2 |
| [4Fe 4S] models |  |  |  |  |  |  |  |  |  |
| 4S 3Fe | 4 | 2.259 (18) | 6.5 (10) | - | - | 2.74 (3) | 13 (4) | -3 (2) | 7.0 |
| 3.75S<br>0.25O/N<br>3Fe | 3 | 2.260 (17) | 6.2 (8) | 2.05<sup>c</sup> | 5<sup>c</sup> | 2.74 (2) | 13 (3) | -3 (2) | 5.3 |
| 3.5S<br>0.5O/N<br>3Fe | 2 | 2.256 (12) | 5 (3) | 2.2 (4) | 5<sup>c</sup> | 2.74 (3) | 13 (2) | -3(3) | 3.3 |
| 3.25S<br>0.75O/N<br>3Fe | 1 | 2.254 (14) | 5 (2) | 2.15<br>(16) | 5<sup>c</sup> | 2.74 (3) | 13 (2) | -3 (2) | 2.9 |
| 3S 1O/N 3Fe | 0 | 2.262 (13) | 4.5 (10) | 2.08 (7) | 15 (20) | 2.744 (2) | 13 (2) | -1 (2) | 2.4 |

<sup>a</sup>Fixed parameters: numbers of S, O/N, and Fe atoms. The Debye-Waller factors were fitted starting from values comparable with the ones found in the literature for different [Fe-S] clusters: $\sigma^2(\text{Fe-S}) = 0.003 \text{ \AA}^2$ , $\sigma^2(\text{Fe-Fe}) = 0.003 \text{ \AA}^2$ , $\sigma^2(\text{Fe-O}) = 0.005 \text{ \AA}^2$ . <sup>b</sup>R, interatomic distance; $\sigma^2$ , Debye Waller factor; $\Delta E_0$ , shift in the energy origin; $R_{\text{fit}}$ , goodness-of-fit calculated as $\Sigma(x_{\text{exp}} - x_{\text{fit}})^2 / \Sigma(x_{\text{exp}})^2$ , where $x_{\text{exp}}$ is the experimental data point, and $x_{\text{fit}}$ the corresponding point in the best-fitting curve. The error on the last digit is reported in parenthesis. <sup>c</sup>This value was fixed during the fit. In bold are the favored models.

Three models provide the best agreement with the experimental data, corresponding to a goodness-of-fit R=0.6% (Table 1). The first one is 3S 1N/O 1Fe, corresponding to a [2Fe-2S] cluster where the average Fe coordination sphere is composed by 3S and 1N/O atoms (see Figure S2). Considering that 2 sulfur atoms belong to the inorganic sulfides bridging the metal centers, the cluster is anchored to the protein scaffold through two S and two N/O donors. The two other models, 3S 1N/O 1.33Fe and 2.67S 1.33N/O 1.33Fe correspond to linear [3Fe-4S] anchored to the protein through one S and three O/N donors, or through N/O donors only, respectively (see Figure S2). The number of Fe neighbors estimated in the initial fit, in which all parameters were allowed to vary, was 1.3 ± 0.3, consistent with [2Fe-2S] and/or linear [3Fe-4S] cluster(s). Such geometries of Fe-S clusters are in agreement with UV-visible analyses.

### EPR and Mössbauer spectroscopies

In order to confirm the presence of [2Fe-2S] and/or linear [3Fe-4S] clusters within the as-isolated SufBC_2_D complex we performed EPR and Mössbauer spectroscopy on as-isolated SufBC_2_D after expression in the presence of ^57^Fe in culture medium and purification under anaerobic conditions. EPR and Mössbauer spectroscopy on two samples returned identical results. After purification, the ^57^FeS-SufBC_2_D complex contains identical iron and sulfur content (average of 2.4 Fe and 2.3 S/ complex) and UV-vis. spectrum to that of ^56^FeS-SufBC_2_D complex (Figure 1B).

No intense EPR signal was detected for the FeS-SufBC_2_D complex, whatever the conditions used (low/high temperature, power, modes (parallel vs perpendicular) (Figure S3). In perpendicular mode, the low-field spectrum recorded at low temperature (5 K) and high power (10 mW) consisted only of a very small signal at g=4.3, which likely corresponds to adventitiously-bound high-spin (S=5/2) Fe^3+^ species. No signal was visible at lower fields (Figure S3A), in particular around g=9.6, where the spectrum of linear [3Fe-4S]^1+^ usually has additional features.^16, 18^ Based on this result, no such species is present in the sample. Once again in perpendicular mode, but at higher temperature (20K) and lower power (1 mW), accumulation allowed us to observe a signal at g=2.01 (Figure S3B), that may correspond to cubane [3Fe-4S]^1+^ cluster. The very low intensity of this signal made further characterization impossible. In parallel mode, no signal was detected, even after extensive temperature/power optimization (Figure S3C). Taken together, these EPR spectroscopy results suggest a species with diamagnetic or integer spin (lack of signal in parallel mode low-frequency EPR cannot be taken as evidence of lack of integer spin). Alternatively, the signal may be very broad, making its signal-to-noise ratio very low. Attempts to reduce EPR sample with one equivalent of electron vs the protein failed.

Mössbauer spectrum, recorded at high temperature ca. 230 K in a 60 mT external magnetic field applied parallel to the gamma rays, is a superposition of two quadrupole doublets (Figure 3). The parameters of the main doublet (89%, *δ* = 0.23 mm/s, Δ*E*_Q_ = 0.52 mm/s) are consistent with Fe^3+^ ions (S=5/2) in a tetrahedral geometry (Figure 3, in blue). The minority doublet (11%, *δ* = 0.48 mm/s, Δ*E*_Q_ = 1.00 mm/s) has an isomer shift parameter value that lies between those corresponding to Fe^2+^ and Fe^2.5+^ ions^28^ (Figure 3, in purple). At 6 K in a 60 mT external magnetic field applied parallel to the gamma rays, the Mössbauer spectrum is quite complex with contributions between −2 mm/s and +3 mm/s (Figure S4). This spectrum reveals the presence of two species. The first one accounting for ~40% of total iron is characteristic of a [2Fe-2S]^2+^ cluster whose parameters (*δ* = 0.30 mm/s, Δ*E*_Q_ = 0.63 mm/s) are indicative of sulfur and nitrogen coordination. The remaining species (~60%) exhibits magnetic hyperfine structures whose origin is hard to define due to the lack of resolved absorption lines. There is no quantifiable contribution of linear [3Fe-4S]^1+^ whose Mössbauer spectrum would extend from −5 mm/s to +6 mm/s and would display several hyperfine structures (Figure S5A).^16, 18^ In addition, the Mössbauer spectrum recorded at 6 K with a 4 T magnetic field applied parallel to the γ-rays is lacking the intense absorption lines at −4.5 and +5 mm s^−1^ usually observed for linear [3Fe-4S]^1+^ (Figure S5B).^16, 18^ Both Mössbauer spectra also lead to exclude the presence of a cuboidal [3Fe-4S]^1+^ (Figure S5C and 5D), consistently with EPR studies.^29^ Concerning [3Fe-4S], the one-electron reduced form only exists for the cubane structure and all investigated [3Fe-4S]^0^ clusters present a S=2 ground state.^30–33^ The recalculated Mössbauer spectra don’t match the experimental ones (Figure S6), precluding the presence of such a cluster, in agreement with EPR spectra where no feature is detected below 100 mT (Figure S3).^32^ Altogether, these analyses allow us to conclude that as-purified SufBC_2_D complex contains two types of Fe-S clusters, a [2Fe-2S]^2+^ cluster (~40%) and an as yet undescribed new magnetic species (~60%).

**Figure 3:**
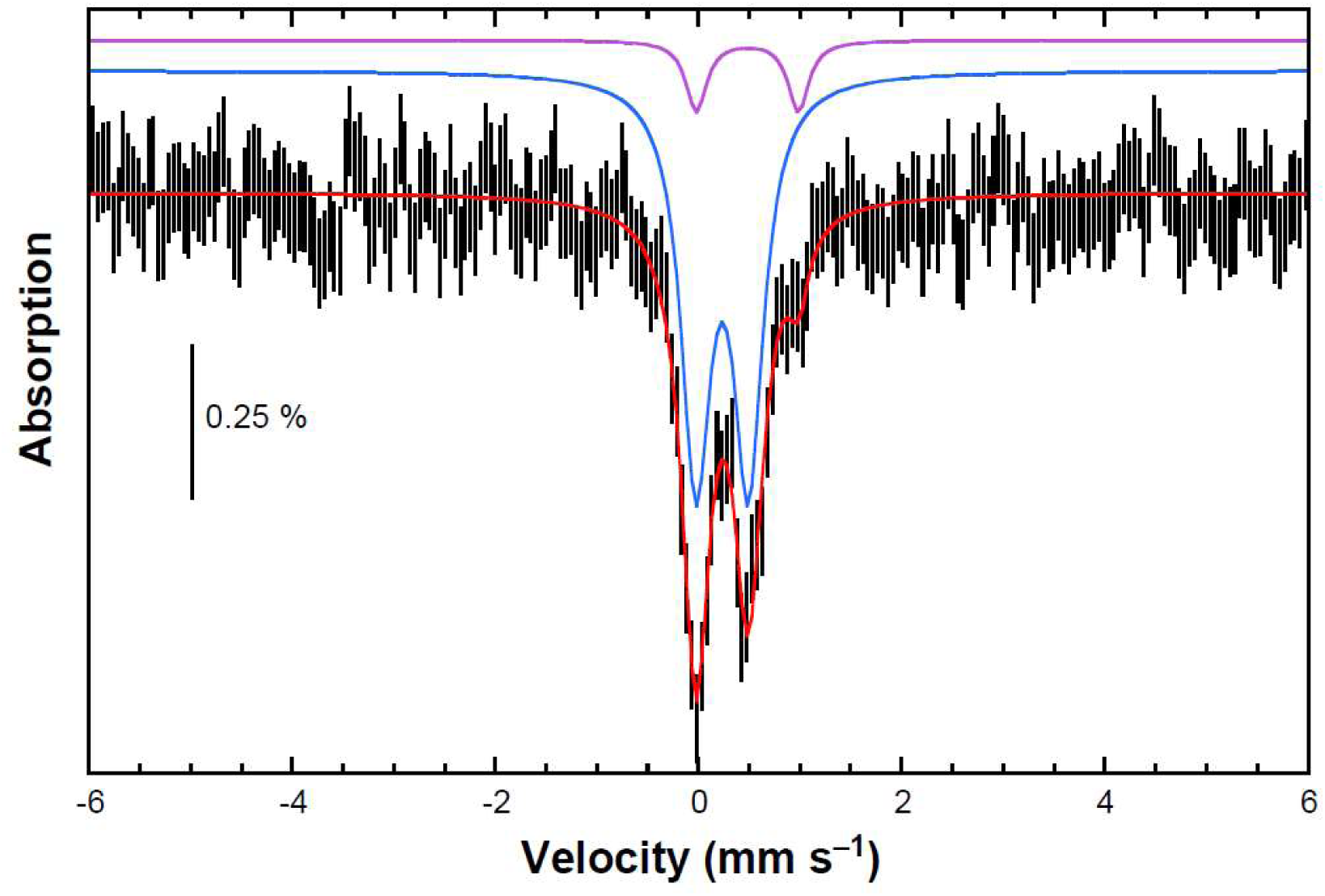
Experimental Mössbauer spectrum of SufBC_2_D complex (290 µM, 2.6 Fe/complex) recorded at ca 230 K with a 0.06 T external magnetic field applied parallel to the γ-rays direction (vertical bars). A simulation is overlaid (solid red line) and the two contributions displayed above as colored traces. See text for parameters.

Taking into account the XAS results, including best fitting models and number of Fe neighbor to the absorber (1.3 +/−0.3), the Fe and S contents per complex after anaerobic purification that is always above 2, and both EPR and Mössbauer analyses, we suggest that this new magnetic species is formed by a new type of “trinuclear” Fe-S system, i.e. a Fe(III) Fe(III) system perturbed by a third probably Fe(II) (to be compatible with EPR). This could correspond to a [3Fe-3S] cluster, a type of Fe-S cluster never reported in the Fe-S biogenesis field.

### Mutagenesis analysis of the SufBC_2_D complex

A previous study carried out *in vivo* proposed SufB Cys405, Glu432, His433 and Glu434 residues, as well as SufD Cys358 and His360 residues as critical residues for Fe-S formation and/or coordination.^15^ Surprisingly, Glu434 and Glu432 point in the opposite direction of the other residues critical for Fe-S assembly (Cys405, His433, Cys358 and His360) (Figure S7). Glu434 is a conserved amino-acid in SufB proteins while Glu432 is not (Figure S8). Cys358 in SufD proteins is not strictly conserved either (Figure S8), but it is one of the mercuric ion ligand in the Hg^2+^-containing SufBC_2_D structure (PDB: 5AWG).^10^ Therefore, we constructed SufB^C405A^CD, SufB^H433A^CD, SufB^E434A^CD, SufBCD^C358A^ and SufBCD^H360A^ variants and assessed their role in cluster binding by analyzing UV-vis. absorption spectroscopy and iron/sulfur content of anaerobically purified variant complexes. With the exception of SufB^E434A^CD variant (Figure S9A-B), SDS-PAGE of variant complexes show that they contain the three proteins SufB, SufC and SufD, as it is observed for the wild-type complex (Figure 1A). Whereas these four mutations had little or no effect on complex formation, distinctive UV-vis. signatures were observed (Figure 1C-1F). Moreover, as detailed below, the Fe and S contents were all significantly lower than those of the wild-type (2.4 Fe and 2.3 S/complex) (Figure 1B). The UV-visible spectra of the SufB^C405A^C_2_D and SufB^H433A^C_2_D variants display small, if any, absorption at 420 nm, and they retain very poor iron binding capacity (0.1 Fe and 0.1 S/complex) (Figure 1C and 1F). The SufBC_2_D^H360A^ variant is also affected in its Fe-S assembly as shown by the low absorption bands in the 320-500 nm region, and the low Fe and S contents, 0.4 Fe and 0.3 S/complex (Figure 1D). Although the UV-vis. spectrum of SufBC_2_D^C358A^ displays low absorption bands, the iron and sulfur content of this complex (1.3 Fe and 1.5 S/complex) suggests that this variant is still able to bind Fe-S cluster (Figure 1E). Surprisingly, as mentioned above, we were unable to isolate the SufB^E434A^C_2_D complex suggesting, in contrast to a previous report, that the mutation E434A in SufB causes a drastic destabilizing effect (Figure S9A-B). In contrast, we were able to purify the SufB^E434K^C_2_D variant. It displays low absorption bands on the UV-vis. absorption spectrum (Figure S9C) and poor levels of iron and sulfur (0.3 Fe and S/complex), indicating that this Glu434 residue is important for Fe-S assembly. Altogether, these data obtained on purified complexes strongly suggest that Cys405, His433 and Glu434 from SufB as well as Cys358 and His360 from SufD are involved in Fe-S cluster ligation/assembly of the *E. coli* SufBC_2_D complex.

### Models for Fe-S clusters on SufBC_2_D

Both spectroscopic data and mutagenesis studies described above provide us with a new and original vision of the Fe-S bound to the SufBC_2_D complex. We propose SufBC_2_D complex to arise under two possible types. One type that accounts for ~40% of the SufBC_2_D complex population would bind a [2Fe-2S] cluster while another type that accounts for ~60% would bind a [3Fe-3S] cluster. We propose these two populations to coexist, at least in the conditions used in our study. We propose the [2Fe-2S] cluster to be coordinated by SufB residues Cys405 and His433 and SufD residues Cys358 and His360. The sulfur coordination provided by Cys405 and Cys358 is consistent with the crystallographic structure of Hg^2+^-containing SufBC_2_D complex in which the two Hg^2+^ ions are bound to Cys405 (in SufB) and Cys358 (in SufD) (PDB: 5AWG).^10^ Cys405 residue is strictly conserved throughout SufB proteins in prokaryotes, including both eubacteria, archaea and plants (Figure S8). In contrast, SufD Cys358 is not conserved in firmicutes such as *Bacillus subtilis* (Figure S8). However, the *E. coli* and *B. subtilis* SUF system show a series of difference from genetic organization to genetic composition and were recently shown to exhibit different efficiency in maturing heterologous Fe-S targets^34, 35^. In contrast, both residues SufB His433 and SufD His360 which we propose to serve as nitrogen ligands are strictly conserved (Figure S8). Moreover, we propose SufB His433 and SufD His360 to coordinate each a different Fe atom of the [2Fe-2S] cluster. The isomer shift which was obtained for the [2Fe-2S] cluster of SufBC_2_D is closer to that observed for ferric sites with 3 Cys and 1 His, as observed for one of the iron site in IscR^36^ or MitoNEET^37^, rather than to that determined in Rieske type proteins^38^ or Apd1 protein^39^ where one iron site is coordinated by 2 His. Thus, we propose that the coordination mode of the [2Fe-2S] cluster in SufBC_2_D is symmetric with two residues from SufB, namely Cys405 and His433, and two residues from SufD, namely Cys358 and His360. To our knowledge, such a coordination mode of a [2Fe-2S] cluster has never been reported. The [3Fe-3S] cluster, like the [2Fe-2S] cluster discussed above, would be coordinated by SufB residues Cys405 and His433 and SufD residue His360, but a striking difference we propose is that an additional inorganic sulfide, instead of Cys358, would bridge a third Fe. This would account for the fact that the SufBC_2_D^C358A^ variant has some Fe-S cluster bound (around 50%). Moreover, the [3Fe-3S] cluster would also be coordinated by SufB Glu434 (bidentate ligand) residue. A water molecule might be the fourth ligand of the third Fe to meet with iron tetrahedral geometry constraints (Figure 4). Glu434 was already reported to be important for Fe-S assembly in *in vivo* experiments^15^ and its implication was confirmed in the present analysis of a purified variant complex. In contrast, the hypothesis of Glu434 acting as a ligand of the [2Fe-2S] appears very unlikely as its side chain is positioned in an opposite direction from the predicted ligands of the [2Fe-2S] cluster (Figure S7). It is worth noting that a sulfide bridging the “2Fe-2S” entity and the third Fe, is reminiscent to what was already observed for HydG.^40^

**Figure 4:**
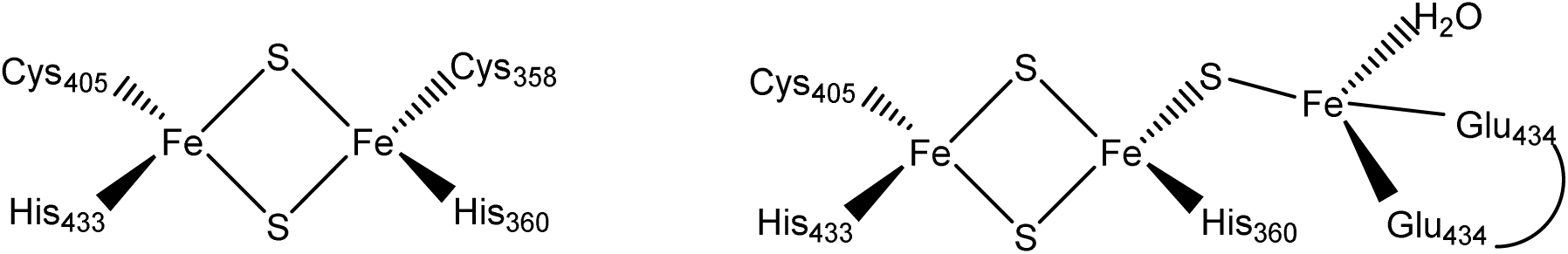
Proposed models of the Fe-S clusters of SufBC_2_D and their ligands.

Finally, it is extremely important to notice that the existence of the two proposed clusters (Figure 4) in the given fractions result in an average Fe coordination sphere composed by 2.6 S and 1.4 O/N atoms and an average of 1.2 second shell Fe, compatible with the values of 2.8 ± 0.3 S, 1.2 ± 0.3 N/O and 1.3 ± 0.3 Fe, extracted from the initial EXAFS analysis.

The current admitted view is that bacterial IscU and SufBC_2_D act as scaffolds assembling a [2Fe-2S] cluster that is subsequently transferred to A-type proteins, which are thought to convert them to [4Fe-4S] (via a reductive coupling) for targeting to apo-targets.^41–47^ Note however that A-type are also instrumental in maturation of [2Fe-2S] containing proteins such as IscR or SoxR in *E. coli*.^48, 49^ The observation of a [2Fe-2S] cluster on as-purified SufBC_2_D is in agreement with this view. The proposed [3Fe-3S] cluster is reproducibly obtained and therefore is unlikely an artifact. The question arises as whether this cluster bound to the complex has any physiological relevance and arises at all *in vivo*. Considering that A-type carrier proteins (ATC) (SufA, IscA) have also been suggested to bind Fe only^43, 50, 51^ an original thought would be that the [3Fe-3S] bound SufBC_2_D species would combine with Fe-bound ATC towards formation of [4Fe-4S] cluster. Even though we don’t favor such an hypothesis, one cannot exclude that the [3Fe-3S] can be a degradation product of higher nuclear Fe-S species such as [4Fe-4S].

### Conclusion

In this work, we have presented the first XAS spectrum of the anaerobically purified SufBC_2_D scaffold from *E. coli*, as well as its Mössbauer, UV-vis. and EPR analysis. These spectroscopic techniques allowed us to identify two new types of Fe-S clusters, a [2Fe-2S] with an atypical coordination and a cluster with a new nuclearity that we proposed to be a [3Fe-3S]. The detailed biochemical characterization on purified SufBC_2_D wt and variants allowed us to propose residues from SufB and SufD acting as ligands for both types of Fe-S clusters while we argue that inorganic sulfide is also involved in the [3Fe-3S] core. These results demonstrate for the first time that as-isolated SufBC_2_D complex contain a [2Fe-2S] cluster in agreement with its scaffold function. The evidence of the existence of a second species, never characterized before, is completely fascinating in the context of Fe-S assembly. The next challenges will be to characterize it by X-ray crystallography and assess its role as a precursor in [4Fe-4S] cluster formation. There is no doubt that the existence of this Fe-S species is a real stimulation in the field of Fe-S cluster biogenesis and more broadly in the field of bioinorganic chemistry.

## ASSOCIATED CONTENT

### Supporting information

Materials and methods; Additional results of experiments (SEC-MALS-RI, EPR and Mössbauer spectra of SufBC^2^D complex, Biochemical properties of SufB^E434A^C^2^D and SufB^E434K^C^2^D variants, Sequence alignments of SufB and SufD proteins); Graphical representation of the Fe-S clusters associated with models used to fit EXAFS spectrum; Structure (zoom) of the apo-SufBC^2^D complex (PDB 5AWF).

## AUTHOR INFORMATION

### Notes

The Authors declare no competing financial interest.

## ACKNOWLEDGMENT

This work was supported by grants from Agence Nationale Recherche (ANR) ANR FeStreS (ANR-11-BSV3-022-01), the LabEx ARCANE (ANR-11-LABX-0003-01) and the CBH-EUR-GS (ANR-17-EURE-0003). We acknowledge the European Synchrotron Radiation Facility for provision of synchrotron radiation facilities (project LS-2814 on beamline BM30).

## ABBREVIATIONS

Fe-S: Iron-sulfur
EPR: Electronic Paramagnetic Resonance
XAS: X-ray absorption spectroscopy

## SYNOPSIS

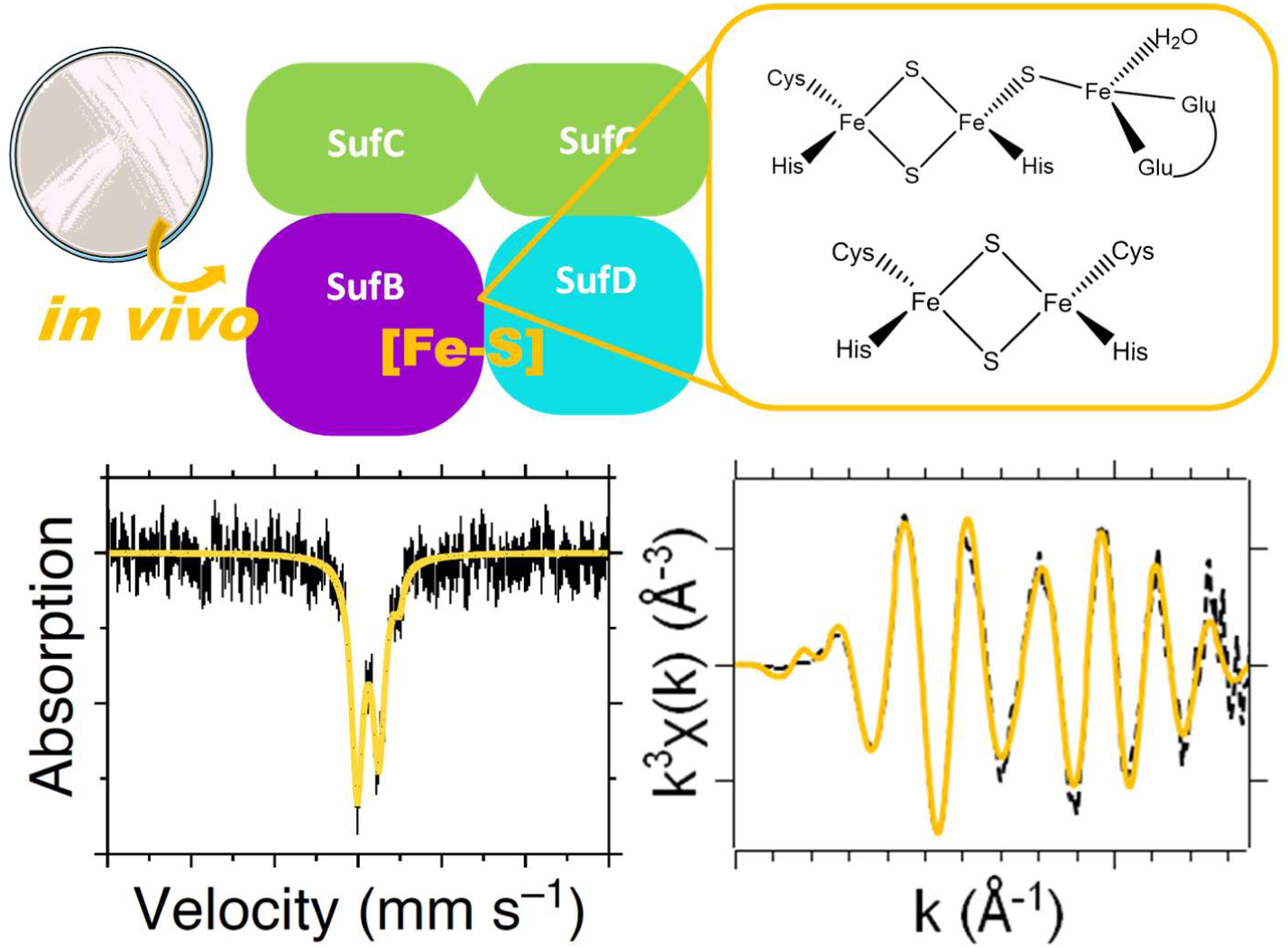

## REFERENCES

1. Fuss, J. O.; Tsai, C. L.; Ishida, J. P.; Tainer, J. A., Emerging critical roles of Fe-S clusters in DNA replication and repair. Biochim. Biophys. Acta 2015, 1853 (6), 1253–1271.

2. Roche, B.; Aussel, L.; Ezraty, B.; Mandin, P.; Py, B.; Barras, F., Iron/sulfur proteins biogenesis in prokaryotes: formation, regulation and diversity. Biochim. Biophys. Acta 2013, 1827 (3), 455–469.

3. Braymer, J. J.; Lill, R., Iron-sulfur cluster biogenesis and trafficking in mitochondria. J. Biol. Chem. 2017, 292 (31), 12754–12763.

4. Roche, B.; Aussel, L.; Ezraty, B.; Mandin, P.; Py, B.; Barras, F., Reprint of: Iron/sulfur proteins biogenesis in prokaryotes: formation, regulation and diversity. Biochim. Biophys. Acta 2013, 1827 (8-9), 923–37.

5. Wollers, S.; Layer, G.; Garcia-Serres, R.; Signor, L.; Clemancey, M.; Latour, J. M.; Fontecave, M.; Ollagnier de Choudens, S., Iron-sulfur (Fe-S) cluster assembly: the SufBCD complex is a new type of Fe-S scaffold with a flavin redox cofactor. J. Biol. Chem. 2010, 285 (30), 23331–23341.

6. Outten, F. W.; Wood, M. J.; Munoz, F. M.; Storz, G., The SufE protein and the SufBCD complex enhance SufS cysteine desulfurase activity as part of a sulfur transfer pathway for Fe-S cluster assembly in Escherichia coli. J. Biol. Chem. 2003, 278 (46), 45713–45719.

7. Loiseau, L.; Ollagnier-de-Choudens, S.; Nachin, L.; Fontecave, M.; Barras, F., Biogenesis of Fe-S cluster by the bacterial Suf system: SufS and SufE form a new type of cysteine desulfurase. J. Biol. Chem. 2003, 278 (40), 38352–38359.

8. Gupta, V.; Sendra, M.; Naik, S. G.; Chahal, H. K.; Huynh, B. H.; Outten, F. W.; Fontecave, M.; Ollagnier de Choudens, S., Native Escherichia coli SufA, coexpressed with SufBCDSE, purifies as a [2Fe-2S] protein and acts as an Fe-S transporter to Fe-S target enzymes. J. Am. Chem. Soc. 2009, 131 (17), 6149–6153.

9. Chahal, H. K.; Outten, F. W., Separate FeS scaffold and carrier functions for SufB(2)C(2) and SufA during in vitro maturation of [2Fe2S] Fdx. J. Inorg. Biochem. 2012, 116, 126–134.

10. Hirabayashi, K.; Yuda, E.; Tanaka, N.; Katayama, S.; Iwasaki, K.; Matsumoto, T.; Kurisu, G.; Outten, F. W.; Fukuyama, K.; Takahashi, Y.; Wada, K., Functional Dynamics Revealed by the Structure of the SufBCD Complex, a Novel ATP-binding Cassette (ABC) Protein That Serves as a Scaffold for Iron-Sulfur Cluster Biogenesis. J. Biol. Chem. 2015, 290 (50), 29717–29731.

11. Saini, A.; Mapolelo, D. T.; Chahal, H. K.; Johnson, M. K.; Outten, F. W., SufD and SufC ATPase activity are required for iron acquisition during in vivo Fe-S cluster formation on SufB. Biochemistry 2010, 49 (43), 9402–9412.

12. Chahal, H. K.; Dai, Y.; Saini, A.; Ayala-Castro, C.; Outten, F. W., The SufBCD Fe-S scaffold complex interacts with SufA for Fe-S cluster transfer. Biochemistry 2009, 48 (44), 10644–10653.

13. Layer, G.; Gaddam, S. A.; Ayala-Castro, C. N.; Ollagnier-de Choudens, S.; Lascoux, D.; Fontecave, M.; Outten, F. W., SufE transfers sulfur from SufS to SufB for iron-sulfur cluster assembly. J. Biol. Chem. 2007, 282 (18), 13342–13350.

14. Blanc, B.; Clemancey, M.; Latour, J. M.; Fontecave, M.; Ollagnier de Choudens, S., Molecular investigation of iron-sulfur cluster assembly scaffolds under stress. Biochemistry 2014, 53 (50), 7867–7869.

15. Yuda, E.; Tanaka, N.; Fujishiro, T.; Yokoyama, N.; Hirabayashi, K.; Fukuyama, K.; Wada, K.; Takahashi, Y., Mapping the key residues of SufB and SufD essential for biosynthesis of iron-sulfur clusters. Sci. Rep. 2017, 7 (1), 9387.

16. Kennedy, M. C.; Kent, T. A.; Emptage, M.; Merkle, H.; Beinert, H.; Munck, E., Evidence for the formation of a linear [3Fe-4S] cluster in partially unfolded aconitase. J. Biol. Chem. 1984, 259 (23), 14463–14471.

17. Jones, K.; Gomes, C. M.; Huber, H.; Teixeira, M.; Wittung-Stafshede, P., Formation of a linear [3Fe-4S] cluster in a seven-iron ferredoxin triggered by polypeptide unfolding. J. Biol. Inorg. Chem. 2002, 7 (4-5), 357–362.

18. Zhang, B.; Bandyopadhyay, S.; Shakamuri, P.; Naik, S. G.; Huynh, B. H.; Couturier, J.; Rouhier, N.; Johnson, M. K., Monothiol glutaredoxins can bind linear [Fe3S4]+ and [Fe4S4]2+ clusters in addition to [Fe2S2]2+ clusters: spectroscopic characterization and functional implications. J. Am. Chem. Soc. 2013, 135 (40), 15153–15164.

19. Musgrave, K. B.; Angove, H. C.; Burgess, B. K.; Hedman, B.; Hodgson, K. O., All-ferrous titanium(III) citrate reduced Fe protein of nitrogenase: An XAS study of electronic and metrical structure. J. Am. Chem. Soc. 1998, 120 (21), 5325–5326.

20. Kounosu, A.; Li, Z. R.; Cosper, N. J.; Shokes, J. E.; Scott, R. A.; Imai, T.; Urushiyama, A.; Iwasaki, T., Engineering a three-cysteine, one-histidine ligand environment into a new hyperthermophilic archaeal Rieske-type [2Fe-2S] ferredoxin from Sulfolobus solfataricus. J. Biol. Chem. 2004, 279 (13), 12519–12528.

21. Kowalska, J.; DeBeer, S., The role of X-ray spectroscopy in understanding the geometric and electronic structure of nitrogenase. Bba-Mol. Cell. Res. 2015, 1853 (6), 1406–1415.

22. Kowalska, J. K.; Hahn, A. W.; Albers, A.; Schiewer, C. E.; Bjornsson, R.; Lima, F. A.; Meyer, F.; DeBeer, S., X-ray Absorption and Emission Spectroscopic Studies of [L2Fe2S2](n) Model Complexes: Implications for the Experimental Evaluation of Redox States in Iron-Sulfur Clusters. Inorg. Chem. 2016, 55 (9), 4485–4497.

23. Bhave, D. P.; Han, W. G.; Pazicni, S.; Penner-Hahn, J. E.; Carroll, K. S.; Noodleman, L., Geometric and Electrostatic Study of the [4Fe-4S] Cluster of Adenosine-5 ‘-Phosphosulfate Reductase from Broken Symmetry Density Functional Calculations and Extended X-ray Absorption Fine Structure Spectroscopy. Inorg. Chem. 2011, 50 (14), 6610–6625.

24. Hsueh, K. L.; Yu, L. K.; Chen, Y. H.; Cheng, Y. H.; Hsieh, Y. C.; Ke, S. C.; Hung, K. W.; Chen, C. J.; Huang, T. H., FeoC from Klebsiella pneumoniae Contains a [4Fe-4S] Cluster. J. Bacteriol. 2013, 195 (20), 4726–4734.

25. Vaccaro, B. J.; Clarkson, S. M.; Holden, J. F.; Lee, D. W.; Wu, C. H.; Poole, F. L.; Cotelesage, J. J. H.; Hackett, M. J.; Mohebbi, S.; Sun, J. C.; Li, H. L.; Johnson, M. K.; George, G. N.; Adams, M. W. W., Biological iron-sulfur storage in a thioferrate-protein nanoparticle. Nat. Commun. 2017, 8, 16110.

26. Cosper, N. J.; Eby, D. M.; Kounosu, A.; Kurosawa, N.; Neidle, E. L.; Kurtz, D. M.; Iwasaki, T.; Scott, R. A., Redox-dependent structural changes in archaeal and bacterial Rieske-type [Ne-2S] clusters. Prot. Sci. 2002, 11 (12), 2969–2973.

27. Blank, M. A.; Lee, C. C.; Hu, Y.; Hodgson, K. O.; Hedman, B.; Ribbe, M. W., Structural models of the [Fe4S4] clusters of homologous nitrogenase Fe proteins. Inorg. Chem. 2011, 50 (15), 7123–7128.

28. Wu, C. H.; Jiang, W.; Krebs, C.; Stubbe, J., YfaE, a ferredoxin involved in diferric-tyrosyl radical maintenance in Escherichia coli ribonucleotide reductase. Biochemistry 2007, 46 (41), 11577–11588.

29. Krebs, C.; Henshaw, T. F.; Cheek, J.; Huynh, B. H.; Broderick, J. B., Conversion of 3Fe-4S to 4Fe-4S clusters in native pyruvate formate-lyase activating enzyme: Mossbauer characterization and implications for mechanism. J. Am. Chem. Soc. 2000, 122 (50), 12497–12506.

30. Zhang, B.; Arcinas, A. J.; Radle, M. I.; Silakov, A.; Booker, S. J.; Krebs, C., First Step in Catalysis of the Radical S-Adenosylmethionine Methylthiotransferase MiaB Yields an Intermediate with a [3Fe-4S](0)-Like Auxiliary Cluster. J. Am. Chem. Soc. 2020, 142 (4), 1911–1924.

31. Hu, Z.; Jollie, D.; Burgess, B. K.; Stephens, P. J.; Munck, E., Mossbauer and EPR studies of Azotobacter vinelandii ferredoxin I. Biochemistry 1994, 33 (48), 14475–14485.

32. Papaefthymiou, V. G. J. J.; Moura, I.; Moura, J.J.G.; Münck, E., Mössbauer study of D. gigas ferredoxin II and spin-coupling model for the Fe3S4 cluster with valence delocalization. J. Am. Chem. Soc. 1987, 109, 4703–4710.

33. Lanz, N. D. P. M. E.; Kakar, E. S.; Lee, K. H.; Krebs, C.; Booker, S. J., Evidence for a catalytically and kinetically competent enzyme–substrate cross-linked intermediate in catalysis by Lipoyl Synthase. Biochemistry 1994, 53, 4557–4572.

34. Santos, P. C. D. B., subtilis as a Model for Studying the Assembly of Fe–S Clusters in Gram-Positive Bacteria. In Methods in Enzymolology, 2017; Vol. 595, p 186.

35. Francesca D’Angelo, E. F.-F., Pierre Simon Garcia, Helena Shomar, Martin Pelosse, Rita Rebelo Manuel, Ferhat Büke, Siyi Liu, Niels van den Broek, Nicolas Duraffourg, Carol de Ram, Martin Pabst, Emmanuelle Bouveret, Simonetta Gribaldo, Béatrice Py, Sandrine Ollagnier de Choudens, Frédéric Barras, Gregory Bokinsky, Cellular assays identify barriers impeding iron-sulfur enzyme activity in a non-native prokaryotic host. eLIFE 2022 (in press).

36. Fleischhacker, A. S.; Stubna, A.; Hsueh, K. L.; Guo, Y.; Teter, S. J.; Rose, J. C.; Brunold, T. C.; Markley, J. L.; Munck, E.; Kiley, P. J., Characterization of the [2Fe-2S] cluster of Escherichia coli transcription factor IscR. Biochemistry 2012, 51 (22), 4453–4462.

37. Ferecatu, I.; Goncalves, S.; Golinelli-Cohen, M. P.; Clemancey, M.; Martelli, A.; Riquier, S.; Guittet, E.; Latour, J. M.; Puccio, H.; Drapier, J. C.; Lescop, E.; Bouton, C., The diabetes drug target MitoNEET governs a novel trafficking pathway to rebuild an Fe-S cluster into cytosolic aconitase/iron regulatory protein 1. J. Biol. Chem. 2014, 289 (41), 28070–28086.

38. Wolfe, M. D.; Altier, D. J.; Stubna, A.; Popescu, C. V.; Munck, E.; Lipscomb, J. D., Benzoate 1,2-dioxygenase from Pseudomonas putida: single turnover kinetics and regulation of a two-component Rieske dioxygenase. Biochemistry 2002, 41 (30), 9611–9626.

39. Stegmaier, K.; Blinn, C. M.; Bechtel, D. F.; Greth, C.; Auerbach, H.; Muller, C. S.; Jakob, V.; Reijerse, E. J.; Netz, D. J. A.; Schunemann, V.; Pierik, A. J., Apd1 and Aim32 Are Prototypes of Bishistidinyl-Coordinated Non-Rieske [2Fe-2S] Proteins. J. Am. Chem. Soc. 2019, 141 (14), 5753–5765.

40. Dinis, P.; Suess, D. L.; Fox, S. J.; Harmer, J. E.; Driesener, R. C.; De La Paz, L.; Swartz, J. R.; Essex, J. W.; Britt, R. D.; Roach, P. L., X-ray crystallographic and EPR spectroscopic analysis of HydG, a maturase in [FeFe]-hydrogenase H-cluster assembly. Proc. Natl. Acad. Sci. U S A 2015, 112 (5), 1362–1367.

41. Vinella, D.; Brochier-Armanet, C.; Loiseau, L.; Talla, E.; Barras, F., Iron-sulfur (Fe/S) protein biogenesis: phylogenomic and genetic studies of A-type carriers. PLoS Genet. 2009, 5 (5), e1000497.

42. Tan, G.; Lu, J.; Bitoun, J. P.; Huang, H.; Ding, H., IscA/SufA paralogues are required for the [4Fe-4S] cluster assembly in enzymes of multiple physiological pathways in Escherichia coli under aerobic growth conditions. Biochem. J. 2009, 420 (3), 463–472.

43. Wang, W.; Huang, H.; Tan, G.; Si, F.; Liu, M.; Landry, A. P.; Lu, J.; Ding, H., In vivo evidence for the iron-binding activity of an iron-sulfur cluster assembly protein IscA in Escherichia coli. Biochem. J. 2010, 432 (3), 429–436.

44. Hasnat, M. A.; Zupok, A.; Olas, J. J.; Mueller-Roeber, B.; Leimkuhler, S., A-Type Carrier Proteins Are Involved in [4Fe-4S] Cluster Insertion into the Radical S-Adenosylmethionine Protein MoaA for the Synthesis of Active Molybdoenzymes. J. Bacteriol. 2021, 203 (12), e0008621.

45. Beilschmidt, L. K.; Ollagnier de Choudens, S.; Fournier, M.; Sanakis, I.; Hograindleur, M. A.; Clemancey, M.; Blondin, G.; Schmucker, S.; Eisenmann, A.; Weiss, A.; Koebel, P.; Messaddeq, N.; Puccio, H.; Martelli, A., ISCA1 is essential for mitochondrial Fe4S4 biogenesis in vivo. Nat. Commun. 2017, 8, 15124.

46. Muhlenhoff, U.; Richter, N.; Pines, O.; Pierik, A. J.; Lill, R., Specialized function of yeast Isa1 and Isa2 proteins in the maturation of mitochondrial [4Fe-4S] proteins. J. Biol. Chem. 2011, 286 (48), 41205–41216.

47. Weiler, B. D.; Bruck, M. C.; Kothe, I.; Bill, E.; Lill, R.; Muhlenhoff, U., Mitochondrial [4Fe-4S] protein assembly involves reductive [2Fe-2S] cluster fusion on ISCA1-ISCA2 by electron flow from ferredoxin FDX2. Proc. Natl. Acad. Sci. U S A 2020, 117 (34), 20555–20565.

48. Vinella, D.; Loiseau, L.; Ollagnier de Choudens, S.; Fontecave, M.; Barras, F., In vivo [Fe-S] cluster acquisition by IscR and NsrR, two stress regulators in Escherichia coli. Mol. Microbiol. 2013, 87 (3), 493–508.

49. Gerstel, A.; Zamarreno Beas, J.; Duverger, Y.; Bouveret, E.; Barras, F.; Py, B., Oxidative stress antagonizes fluoroquinolone drug sensitivity via the SoxR-SUF Fe-S cluster homeostatic axis. PLoS Genet. 2020, 16 (11), e1009198.

50. Mapolelo, D. T.; Zhang, B.; Naik, S. G.; Huynh, B. H.; Johnson, M. K., Spectroscopic and functional characterization of iron-bound forms of Azotobacter vinelandii (Nif)IscA. Biochemistry 2012, 51 (41), 8056–8070.

51. Sendra, M.; Ollagnier de Choudens, S.; Lascoux, D.; Sanakis, Y.; Fontecave, M., The SUF iron-sulfur cluster biosynthetic machinery: sulfur transfer from the SUFS-SUFE complex to SUFA. FEBS Lett. 2007, 581 (7), 1362–1368.

