## Supplementary material for "A multidisciplinary analysis of the as-isolated *Escherichia coli* SufBC_2_D complex reveals the presence of two new iron-sulfur clusters": SI_veronesi_BioRxiv.pdf

#### Experimental part.

***Plasmids and Strains.*** His$_6$-SufBCD was co-expressed with SufSE from the pETDuet-1 vector. SufS and SufE were amplified by PCR as one DNA fragment using MG1655 genomic DNA as a template, Phusion polymerase-HF (Thermo Fisher) as DNA polymerase and primers 5'-GGGAATTCCATATGATTTTTTCCGTCGACAAAGTGCGGGCCGACTTTCCGGTGC-3' and 5'-GGGAATTCGGTACCTTAGCTAAGTGCAGCGGCTTTGGCGCGAATTGCGCGAATCAT-3'. The PCR product was digested with NdeI and KpnI and cloned into the corresponding MCS2 sites of pETDuet-1 (Novagen), generating plasmid pETDuet1sufSE. SufBCD were amplified by PCR as one DNA fragment using MG1655 genomic DNA as a template, Phusion polymerase-HF (Thermo Fisher) as DNA polymerase and primers 5'-GGGAATTCGAATTCGTCTCGTAATACTGAAGCAACTGACGATGTCAAAAC-3' and 5'-GGGAATTCCTGCAGTCATCTTGCACCTCCTGGCAGCCGTTGACCGATTCG-3'. The PCR product was digested with EcoRI and PstI and cloned into the corresponding MCS1 sites of pETDuet1sufSE generating plasmid pETDuet-1SufBCDSE. The sequences of all plasmid inserts were confirmed by DNA sequencing. For His$_6$-SufBC$_2$D variants, pETDuet-1SufBCDSE vector was submitted to site directed mutagenesis with appropriate primers (Table S1). Briefly, PCR was carried out using Phusion polymerase-HF (Thermo fischer) at the recommended TM. PCR samples were incubated with a reaction buffer containing 2UI of DpnI, 10 UI of T4 DNA Ligase, 1mM of ATP and 2 UI PNK (Polynucleotide kinase) for 45 min at RT before transformation in Top10 competent cells. Each mutant was DNA-sequenced (MWG) before to be transfer in Rosetta 2(DE3) expression cell.

***Protein expression and purification.*** $His_6$-$SufBC_2D$ (wt and variants) were overexpressed in Rosetta 2(DE3) *Escherichia coli* strain with an auto inducible medium (10 g N-Z-amine, 5 g yeast extract, 100 mM $PO_4$, 25 mM $SO_4$, 50 mM $NH_4$, 100 mM Na, 50 mM K, 0.5 % glycerol, 0.05% glucose, 0.2% α-lactose and 1mM of $MgSO_4$) for 3 h at 37°C and 24h at 20°C. For Mössbauer analysis, cells were cultivated in M9 medium at 37°C, and when the optical density at 600 nm reached 0.5, $^{57}FeCl_3$ (100 μM) was added and then IPTG (0.5 mM) to induce protein expression that was pursued for 3 h. All the following steps were done anaerobically in a glove box. The bacterial pellet was resuspended in buffer A (50 mM Tris, 150 mM NaCl , 50 mM R/E (Glutamate/ Arginine) pH 8) and the lysis was achieved by sonication at 4°C in the presence of a protease inhibitors cocktail (complete, EDTA free from Roche) and 2mg of egg lysozyme (Sigma Aldrich). After centrifugation at 75000 g (20 min at 4°C), the soluble fraction was loaded on a TALON affinity column (New England Biolabs) and two washing steps were performed : the first one with buffer B (50 mM Tris, 1M NaCl, 50mM R/E, pH 8) and the second one with buffer A. The $His_6$-$SufBC_2D$ complex was then eluted with buffer C (buffer A with 300 mM imidazole) and concentrated with a 50 kDa molecular weight cut-off Ultracell concentrator (Amicon). The iron and sulfur content of the protein was determined as described previously.[1, 2]

***SEC-MALS-RI.***

Twenty microlitres of sample was loaded on an analytical Superdex S200 increase size exclusion chromatography (SEC) column (GE Healthcare) pre-equilibrated with runnig buffer (50mM tris pH 7.5, 150mM NaCl). SEC was performed at 0.5 ml.min$^{-1}$ with an in-line multi-angle laser light scattering (MALLS) spectrometer (DAWN HELEOS II, Wyatt Instruments). An in-line refractive index (RI) detector (Optirex, Wyatt Instruments) was used to follow the differential refractive index relative to the solvent. Masses were estimated with the Debye model using ASTRA software version 6 (Wyatt Instruments) with a theoretical dn/dc value of 0.185 $ml.g^{-1}$.

***XAS data acquisition and analysis.*** Fe K-edge X-ray Absorption Spectroscopy was performed at the beamline CRG-FAME-BM30B at the European Synchrotron Radiation Facility (ESRF, Grenoble, France).[3] The EXAFS sample was concentrated at 1.4 mM and 10% glycerol was added to the solution as a cryoprotectant. Drops of the solution (~50 µl) were deposited on the sample holder equipped with kapton windows, immediately frozen in liquid nitrogen, then transferred into the He-cryostat and measured at 15 K. The Fe K-edge energy region was scanned in the 7000-8000 eV range with a nitrogen-cooled Si(220) monochromator. The photon energy was calibrated over a metallic Fe foil, by defining the first inflection point of its absorption spectrum at 7112.0 eV. X-ray absorption spectra were recorded in fluorescence mode with a 30-elements Ge solid-state detector (Canberra). Six scans were acquired over different spots on the sample to avoid radiation damage, providing $\sim 10^6$ total counts in the final spectrum. XAS data reduction and normalization were performed with standard methods using the Athena software.[4] Scattering amplitudes and phase shifts for *ab initio* EXAFS fitting were calculated with the code FEFF6[4, 5] implemented in Artemis. The amplitude reduction factor $S_0^2$ was calculated by the FEFF program from atomic overlap integrals: the value of 0.94 was kept constant during the fit. The fits to the experimental data were performed with the Artemis program, minimizing the R-factor (defined as: $\Sigma(x_{exp}-x_{fit})^2/\ \Sigma(x_{exp})^2$, where $x_{exp}$ are the experimental data points, and $x_{fit}$ the corresponding point in the best fitting curve) with Levenberg-Marquardt non-linear least-squares minimization algorithm.[4, 6] The spectra were Fourier Transformed in the k range 2.8-13.5 $Å^{-1}$, then fitted in the real space, in the range [1.0-3.0] Å. Three single scattering paths (Fe-S, Fe-N/O, Fe-Fe) were included in the model. In order to disentangle the correlation between the multiplicity of a path and its Debye-Waller (DW) factor, nineteen different models were fitted to the experimental spectrum, with fixed path multiplicity, and interatomic distances and DW factors as free parameters.

***Mössbauer spectroscopy.*** For Mössbauer spectroscopy, $SufBC_2D$ sample was prepared in the glove box directly after purification of the complex and transferred to a Mössbauer sample holder, and frozen in cooled 2-methyl-butane inside the glove box and kept in liquid nitrogen until the measurement.

Mössbauer spectra were recorded at 230 K and ca. 6 K, either on a low-field Mössbauer spectrometer equipped with a Janis SVT-400 cryostat, or on a strong-field Mössbauer spectrometer equipped with an Oxford Instruments Spectromag 4000 cryostat containing a 8 T split-pair superconducting magnet. Both spectrometers were operated in a constant acceleration mode in transmission geometry. The isomer shifts are referenced against that of a metallic iron foil at room-temperature. The spectra were analyzed with home-made programs[7, 8]. The same programs were used to calculate theoretical spectra for 3Fe-4S clusters using published parameters.

***EPR spectroscopy.*** X-band Continuous Wave Electron Paramagnetic Resonance (CW-EPR) was performed on an EMX Bruker spectrometer with an ER-4116 dual mode cavity operating at ca. 9.35 GHz (parallel mode) or 9.7 GHz (perpendicular mode) and an Oxford Instrument ESR-900 Helium flow cryostat with a magnetic field modulation of 100 kHz for lock-in amplification.

#### Supplementary Figures and Tables.

##### Table S1: list of primers used in this study to generate SufBCD variants

| Primers names | Primers sequences |
|---|---|
| SufB-C405AFor | 5’ TTTCACTCAG**GCT**GACTCAATGCTGATTGGCGCTAATTG 3’ |
| SufB-C405ARev | 5’ TTGCGCGCATTGGTTGCC 3’ |
| SufB-H433AFor | 5’GCAACTGGAA**GCA**GAGGCAACGAC 3’ |
| SufB-H433ARev | 5’GCACTATTGTTACGACAC 3’ |
| SufB-E434KFor | 5’ ACTGGAACAC**AAA**GCAACGACATC 3’ |
| SufB-E434KRev | 5’ TGCGCACTATTGTTACGAC 3’ |
| SufB-E434AFor | 5’ ACTGGAACAC**GCT**GCAACGACATC 3’ |
| SufB-E434ARev | 5’ TGCGCACTATTGTTACGAC 3’ |
| SufD-C358AFor | 5’ TGATGTGAAA**GCG**AGCCACGGCG 3’ |
| SufD-C358ARev | 5’ TCTGCATAGATTTCCAGC 3’ |
| SufD-H360AFor | 5’ GAAATGCAGC**GCG**GGCGCGACGG 3’ |
| SufD-H360ARev | 5’ ACATCATCTGCATAGATTTC 3’ |

Substituted nucleotides are in bold

**Figure S1**

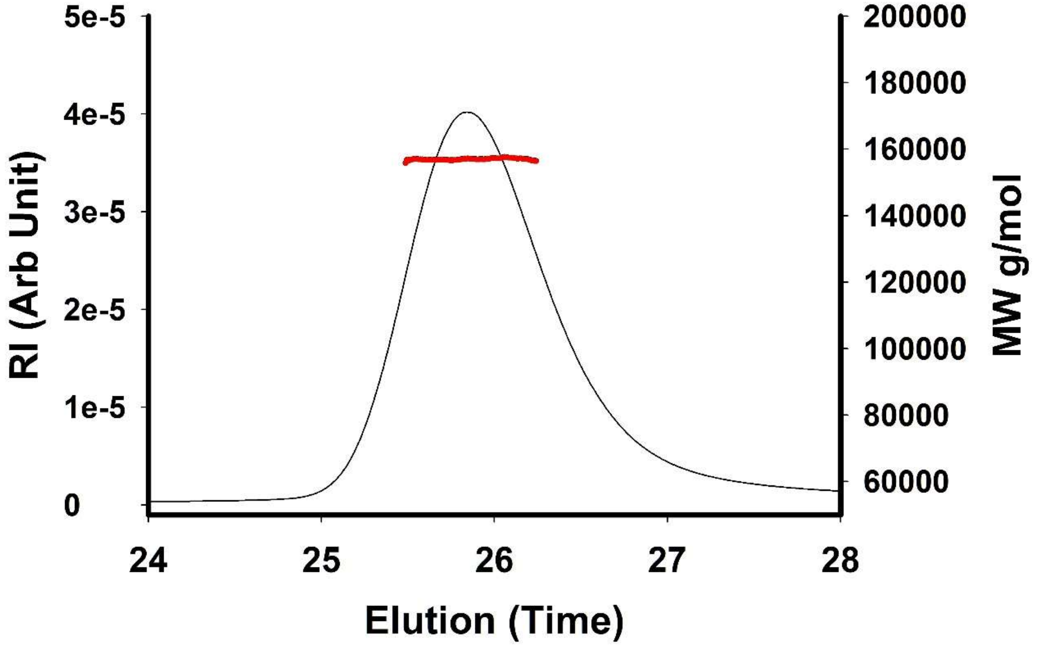


**Figure S1: SEC MALS-RI data of $SufBC_2D$ complex purified under anaerobic conditions**. Sample concentration is determined at 4.8 $mg.ml^{-1}$ and give a $MW_{exp}$ of 157 kDa +/- 0.1%. Differential refractive index RI in Arb Unit (black curve), calculated MW in g/mol (red line).

#### Figure S2

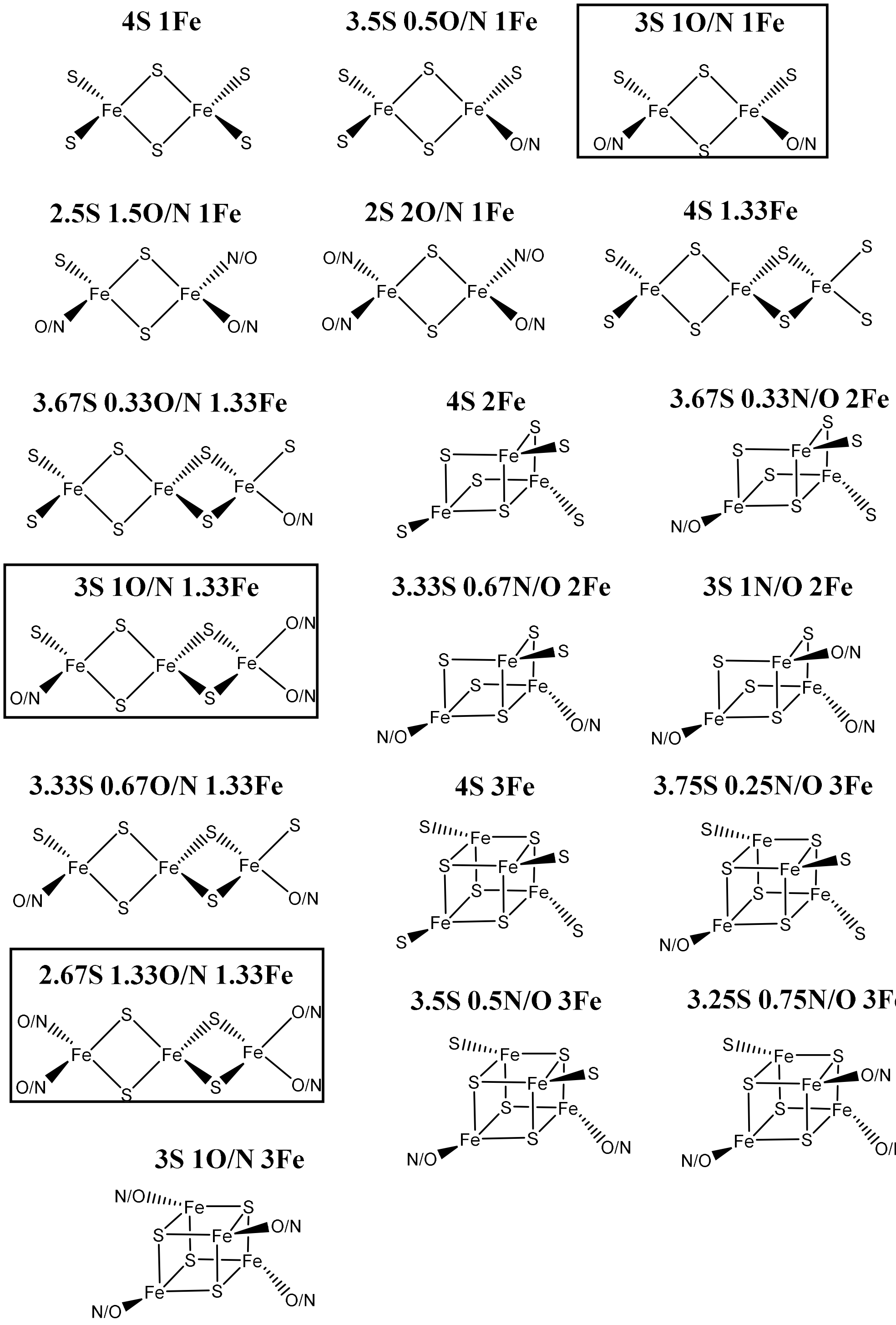


**Figure S2**: **Graphical representation of the [Fe-S] clusters associated with each of the nineteen models used to fit the first two peaks of the Fourier-transformed EXAFS spectrum**. The name of the models (bold) indicate the average number of S and O or N ligands in the coordination sphere of the Fe absorber, followed by the number of second-shell Fe (one in [2Fe-2S], two in [3Fe-4S] and three in [4Fe-4S]). The boxed models correspond to favored models from the fit.

**Figure S3:**

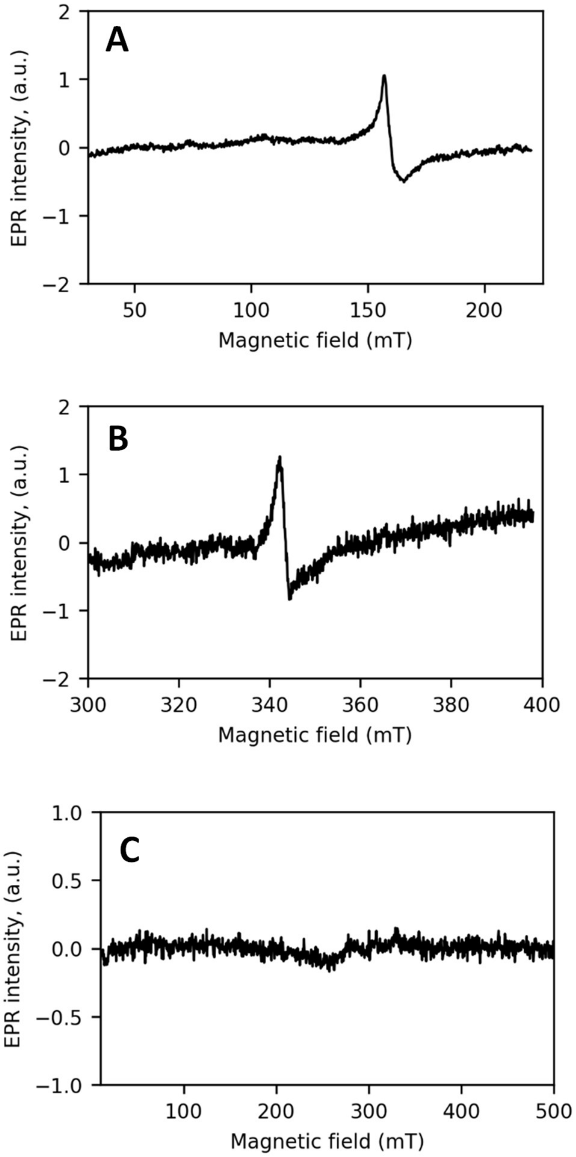


**Figure S3: EPR spectra of SufBC$_2$D complex (150 µM, 400µM iron).** (A): Perpendicular mode, low field and g=4.3 region. T: 5K, P: 10 mW, Scan: 100 ; (B) perpendicular mode, g=2.01 region. T : 20K, P : 1 mW, Scan: 100; (C) Parallel mode. T : 5K, P : 1 mW, Scan: 20.

#### Figure S4:

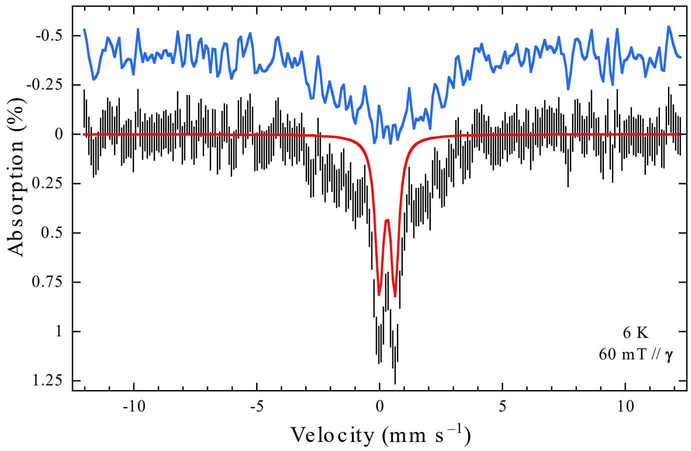


**Figure S4.** Experimental Mössbauer spectrum of $^{57}$Fe-S SufBC$_2$D recorded at ca 6 K with a 0.06 T external magnetic field applied parallel to the γ-beam (vertical bars). The doublet displayed in red accounting for 40% presents the following nuclear parameters: $\delta = 0.30$ mm s$^{-1}$ and $\Delta E_Q = 0.63$ mm s$^{-1}$. The remaining magnetic contribution is shown in blue.

### Figure S5:

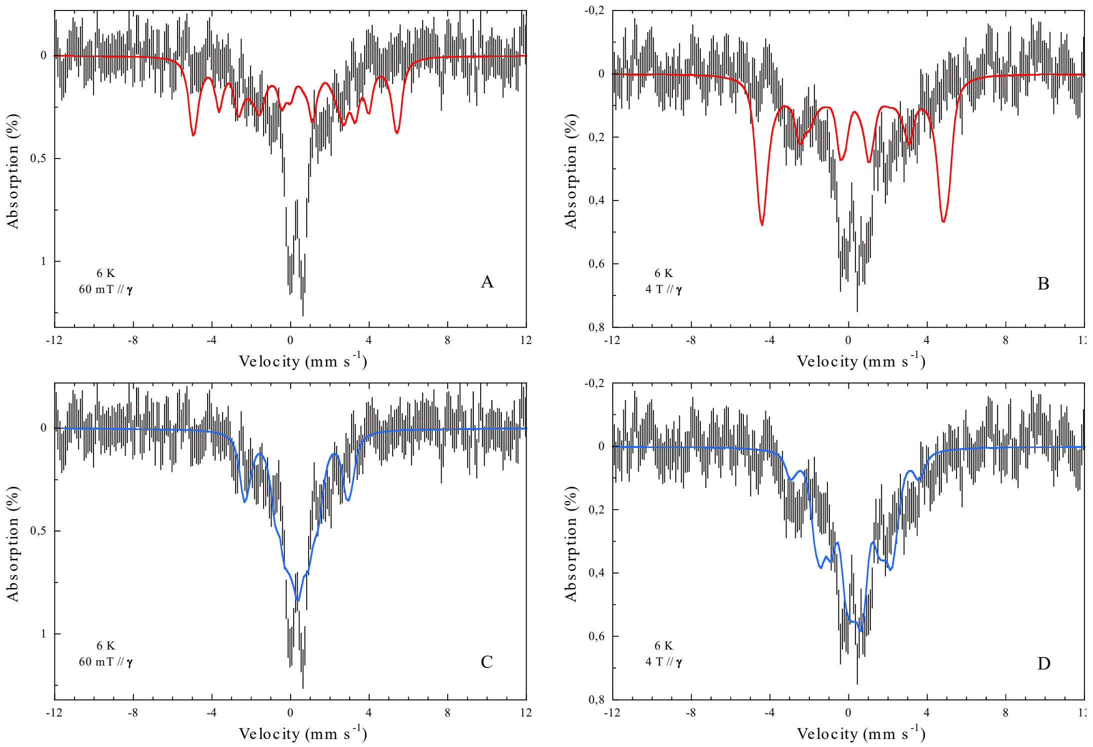


**Figure S5.** Experimental Mössbauer spectra of $^{57}$Fe-S $SufBC_2D$ recorded at ca 6 K with a 0.06 (left column) and 4 T (right column) external magnetic field applied parallel to the γ-beam (vertical bars). Spectra overlaid as solid red lines in panels A and B are associated to the all ferric $[3Fe\text{-}4S]^+$ linear cluster of *Saccharomyces cerevisiae* glutaredoxin 5 (*Sc* Grx5) reconstituted with $^{57}$Fe in the presence of glutathione and were calculated according to the parameters listed in reference[9]. Spectra overlaid as solid blue lines in panels C and D are associated to the all ferric $[3Fe\text{-}4S]^+$ cuboidal cluster in anaerobically purified pyruvate formate-lyase activating enzyme (PFL-AE) and were calculated according to the parameters given in reference[10] (no distribution in exchange constant values was considered). All theoretical spectra were scaled to the area of the experimental spectra.

#### Figure S6

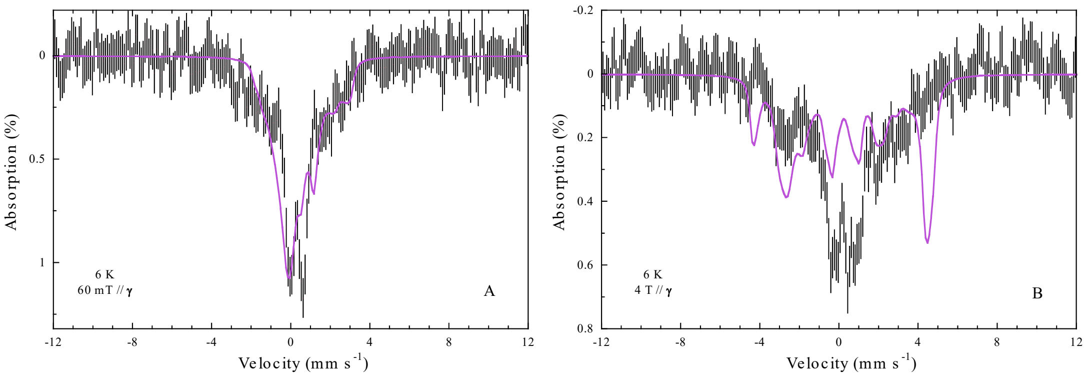


**Figure S6:** Experimental Mössbauer spectra of $^{57}$Fe-S SufBC$_2$D recorded at ca 6 K with a 0.06 (left column) and 4 T (right column) external magnetic field applied parallel to the γ-beam (vertical bars). Spectra overlaid as solid mauve lines in panels A and B are associated to the $[3Fe\text{-}4S]^0$ cuboidal cluster in *Azotobacter vinelandii* ferredoxin I obtained by reduction at pH=8.5 and were calculated according to the parameters given in reference[11]. All theoretical spectra were scaled to the area of the experimental spectra.

**Figure S7**

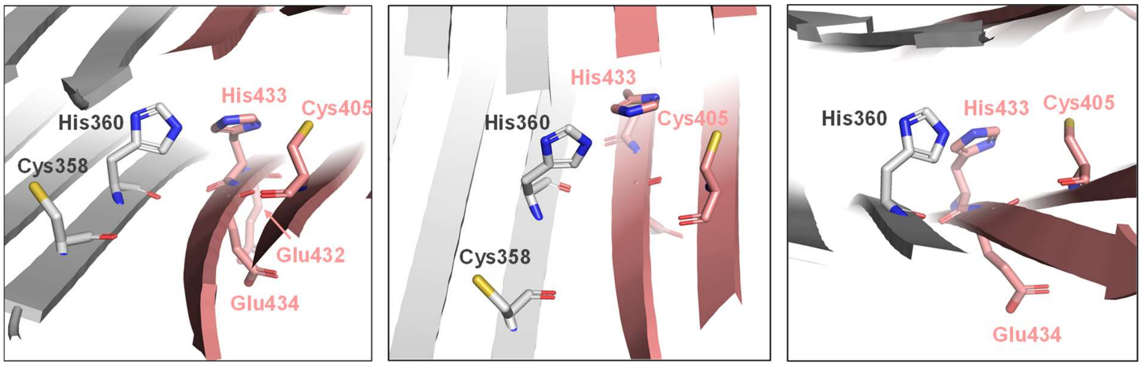


**Figure S7:** Structure (zoom) of the apo-SufBC$_2$D complex (PDB 5AWF) generated with Pymol v2.3.3. Only SufB (light pink) and SufD (grey) are represented. Critical amino-acids (from *in cellulo* and structural analyses) for Fe-S cluster assembly are indicated (left panel); Residues which are proposed in this study to coordinate the [2Fe-2S] cluster (middle panel) and those proposed to ligate the [3Fe-3S] species (right panel).

#### Figure S8: SufB alignment

```
MmSufB       ------------------------------------------------------------ 0
BsSufB       ------------------------------------------------------------ 0
AthSufB      MASLLANGISSFSPQPTSDSSKSPKGFHPKPESLKFPSPKSLNPTRPIFKLRADVGIDSR 60
StSufB       ------------------------------------------------------------ 0
EcSufB       ------------------------------------------------------------ 0
SfSufB       ------------------------------------------------------------ 0


MmSufB       -------------------------------------------------MQTEQVSLKKR 11
BsSufB       --------------------MAK-KMPDIGEYKYGFHDKDVSIFRSERGLTKEIV----E 35
AthSufB      PIGASESSSSGTSTVSSTDKLQQYFQNLDYDKKYGFVE-DIDSFTIPKGLSEETI----R 115
StSufB       -----------MSRNTEATEEVGIWTGGRLNYKEGFFT-QLPTDELAKGISEEVV----R 44
EcSufB       -----------MSRNTEATDDVKTWTGGPLNYKEGFFT-QLATDELAKGINEEVV----R 44
SfSufB       -----------MSRNTEATDDVKTWTGGPLNYKEGFFT-QLATDELAKGINEEVV----R 44
                                                              :  * :    .

MmSufB       AESAAEKKAAFGEDFELEKYEE---------------------GSKVSKPIE-------- 42
BsSufB       EISRMKEEPQWMLDFRLKSLEHFYNMPMPQWGG-DLNSLNFDEITYYVKPSERSER---- 90
AthSufB      LISKLKEEPDWMLEFRFKAYAKFLKLEEPKWSDNRYPSINFQDMCYYSAPKKKPTLNSL- 174
StSufB       AISAKRNEPEWMLEFRLNAYRAWLDMDEPHWLKAHYDKLNYQDYSYYSAPSCGNCDDSCV 104
EcSufB       AISAKRNEPEWMLEFRLNAYRAWLEMEEPHWLKAHYDKLNYQDYSYYSAPSCGNCDDTCA 104
SfSufB       AISAKRNEPEWMLEFRLNAYRAWLEMEEPHWLKAHYDKLNYQDYSYYSAPSCGNCDDTCA 104
               *  .::  :  :*.::                               *

MmSufB       ----------DLQSLDEESKKTLLQVGVIPSEEGRSGSFL---VLDNAV----SHSTLKD 85
BsSufB       ----------SWDEVPEEIKQTFDKLGIPEAEQKYLAGVS--AQYESEVVYHNMKEDLEA 138
AthSufB      ------------DEVDPQLLEYFDKLGVPLTEQKRLANVAVDAVIDSVSIATTHRKTLEK 222
StSufB       SQPGAVQQTGANTFLSKEVEDAFEQLGVPVREGK---EVAVDAIFDSVSVATTYREKLAE 161
EcSufB       SEPGAVQQTGANAFLSKEVEAAFEQLGVPVREGK---EVAVDAIFDSVSVATTYREKLAE 161
SfSufB       SEPGAVQQTGANAFLSKEVEAAFEQLGVPVREGK---EVAVDAIFDSVSVATTYREKLAE 161
                           :  :    :  ::*:   *      .      :.       :. *

MmSufB       KNVELMSTHKAMEKYEW-LKDYSWKLVQVDADKYTAKTYLEDADGYFIRVPAGKKTSMPV 144
BsSufB       QGIVFKDTDSALKENEDIFREHWAKVIPPTDNKFAALNSAVWSGGSFIYVPKGVKVETPL 198
AthSufB      SGVIFCSISEAIREYPDLIKKYLGRVVPSDDNYYAALNSAVFSDGSFCYIPKNTRCPMPI 282
StSufB       QGIIFCSFGEAIHDHPDLVRKYLGTVVPGNDNFFAALNAAVASDGTFIYVPKGVRCPMEL 221
EcSufB       QGIIFCSFGEAIHDHPELVRKYLGTVVPGNDNFFAALNAAVASDGTFIYVPKGVRCPMEL 221
SfSufB       QGIIFCSFGEAIHDHTELVRKYLGTVVPGNDNFFAALNAAVASDGTFIYVPKGVRCPMEL 221
             ..: : .  .*:..    .:.:   ::    : ::* .    :.* *  :* . :    :

MmSufB       QTCLMLGSKKAAQTVHNIIIVEEGATLDIITGCTTKKGVEEGLHLGISEMYIKKGGTLNF 204
BsSufB       QAYFRINSENMGQFERTLIIVDEEASVHYVEGCTAPVYTTNSLHSAVVEIIVKKGGYCRY 258
AthSufB      STYFRINAMETGQFERTLIVAEEGSFVEYLEGCTAPSYDTNQLHAAVVELYCGKGAEIKY 342
StSufB       STYFRINAEKTGQFERTILVADEGSYVSYIEGCSAPVRDSYQLHAAVVEVIIHKDAEVKY 281
EcSufB       STYFRINAEKTGQFERTILVADEDSYVSYIEGCSAPVRDSYQLHAAVVEVIIHKNAEVKY 281
SfSufB       STYFRINAEKTGQFERTILVADEDSYVSYIEGCSAPVRDSYQLHAAVVEVIIHKNAEVKY 281
             .: : :.: : .*  :.:::.:* : :  : **::       ** .: *:   *..  .:

MmSufB       TMIHNWAEQIGVRPRTVVSV------EEGGTYVSNYI-CLKPVRSVQTYPTVRLEGEGAV 257
BsSufB       TTIQNWAN-------NVYNLVTKRTVCEE-NATMEWIDGNIGSKLTMKYPACILKGEGAR 310
AthSufB      STVQNWYAGDEQGKGGIYNFVTKRGLCAGDRSKISWTQVETGSAITWKYPSVVLEGDDSV 402
StSufB       STVQNWFPGDN-NTGGILNFVTKRALCEGENSKMSWTQSETGSAITWKYPSCILRGDNSI 340
EcSufB       STVQNWFPGDN-NTGGILNFVTKRALCEGENSKMSWTQSETGSAITWKYPSCILRGDNSI 340
SfSufB       STVQNWFPGDN-NTGGILNFVTKRALCEGENSKMSWTQSETGSAITWKYPSCILRGDNSI 340
             : ::**          : ..              .:         . .**:  *.*:.:

MmSufB       TRLNTIAIAHPGSELDLGSKAIFNAPGTRAELISRTITIGGRLIARG---EMIGNAKGAK 314
BsSufB       GMTLSIALAGKGQHQDAGAKMIHLAPNTSSTIVSKSISKQGGKVTYRGIVHFGRKAEGAR 370
AthSufB      GEFYSVALTNNYQQADTGTKMIHKGKNTKSRIISKGISAGHSRNCYRGLVQVQSKAEGAK 462
StSufB       GEFYSVALTSGHQQADTGTKMIHIGKNTRSTIISKGISAGHSQNSYRGLVKIMPTATNAR 400
EcSufB       GEFYSVALTSGHQQADTGTKMIHIGKNTKSTIISKGISAGHSQNSYRGLVKIMPTATNAR 400
SfSufB       GEFYSVALTSGHQQADTGTKMIHIGKNTKSTIISKGISAGHSQNSYRGLVKIMPTATNAR 400
                 ::*::   .. * *:* *. . .* : ::*: *:            ..  .* .*:

MmSufB       GHLECKGLVLTDKGSQLAIPILEANVDDIELTHEAAVGKIAKDQVEYLMARGLTEDEAVG 374
BsSufB       SNIECDTLIMDNKSTSDTIPYNEILNDNISLEHEAKVSKVSEEQLFYLMSRGISEEEATE 430
AthSufB      NTSTCDSMLIGDKAAANTYPYIQVKNPSAKVEHEASTSKIGEDQLFYFQQRGIDHERALA 522
StSufB       NFTQCDSMLIGADCGAHTFPYVECRNNSAQLEHEATTSRIGEDQLFYCLQRGISEEDAIS 460
EcSufB       NFTQCDSMLIGANCGAHTFPYVECRNNSAQLEHEATTSRIGEDQLFYCLQRGISEEDAIS 460
SfSufB       NFTQCDSMLIGANCGAHTFPYVECRNNSAQLEHEATTSRIGEDQLFYCLQRGISEEDAIS 460
             .   *. :::  .    : *  :    . .: *** ..::.::*: *   **: .: *

MmSufB       MIIRGFLDVGIRGIPEELKEEIENTIAQTALGM--    407
BsSufB       MIVMGFIEPFTKELPMEYAVEMNRLIKFEMEGSIG    465
AthSufB      AMISGFCRDVFNKLPDEFGAEVNQLMSIKLEGSVG    557
StSufB       MIVNGFCKDVFSELPLEFAVEAQKLLAISLEHSVG    495
```

#### SufD alignment

```
BsSufD          ----------------------------------------------MTLGTKLSVDQEYL   14
SaSufD          -----------------------------------------------MTTDILNISEEQL   13
AthSufD         MAAATVLGRLSLIPNLSSKPKLKSNRRTTSTSVSVRAQASFSDPFVLQLAESLEDSLSAS   60
EcsufD          --------------------------------------------------------MAGL   4
SfSufD          --------------------------------------------------------MAGL   4
StSufD          --------------------------------------------------------MAGL   4


BsSufD          KSFSEKHQEPAWLK----------NLRLQALEQAEDLPMPKPDKTKITNWNFTNFAKHTV   64
SaSufD          VDYSKAHNEPSWMT----------ELRKKALKLTETLEMPKPDKTKLRKWDFDSFKQHDV   63
AthSufD         PSSSLP--LQRI--------------RDSSAETLLSTPWP---SRKDEPFRFTDTSLIRS   101
EcsufD          PNSSNA--LQQWHHLFEAEGTKRSPQAQQHLQQLLRTGLP---TRKHENWKYTPLEGLIN   59
SfSufD          PNSSNA--LQQWHHLFEAEGTKRSPQAQQHLQQLLRTGLP---TRKHENWKYTPLEGLTN   59
StSufD          PNSSKA--LQQWQHLFEEKGESRTEQARQHLQQMLRLGLP---TRKHEDWKYTPLEGLTH   59
                 . *                        .  :       *   . *   : :

BsSufD          DNEPLSSLEDLTDEVKALIDIEN----EDKTLYVQRDQTPAHLSLSQELKDKGVIFT---   117
SaSufD          KGDVYQSLSQLPESVREIIDVDH----SK-NLVIQHNNTIAYTQVDDNASKDGVIVE---   115
AthSufD         SQIE-----PISTQQRNSEILDNLTETQFTNAVIIDG-FVSNLTIGPSDLPDGVYF----   151
EcsufD          SQFVSI-AGEISPQQRDALALTL----DSVRLVFVDGRYVPALS----DATEGSGYEVSI   110
SfSufD          SQFVSI-AGEISPQQRDALALTL----DAVRLVFVDGRYVPALS----DAIEGSGYEVSI   110
StSufD          SQFIQQ-CATISAAQRDALALQI----DAVRLVFVDGRFMPELS----DSTQNSGFDVSV   110
                .         :    :    :      .     .  .              ..

BsSufD          -----DILTAAREHSDLVEKYFMKDGVKVDEHKLTALHAALVNGGAFLYVPKNVQVETPV   172
SaSufD          -----GLADALMNHSDLVQKYFMKDAVTVDEHRITALHTALVNGGVFVYVPKNVVVEHPV   170
AthSufD         -GKYSGLPDELTNRISEFIGNFD------SGDLFWSINGMGAPDLMVIYVPEGCKVENPI   204
EcsufD          NDDRQGLPDAIQ------------------AEVFLHLTESLAQSVTHIAVKRGQRPAKPL   152
SfSufD          NDDRQGLPDAIQ------------------AEVFLHLTESLAQSVTHIAVKRGQRPAKPL   152
StSufD          RDERQALAAPVQ------------------PEVFLHLTESLAQCVTYIQVRRNQRPTRPL   152
                      :                        . :  :    .     : * ..     *:

BsSufD          QAVYVHESN---------DTALFNHVLIVAEDHSSVTYVENYISTVNPKDAVFNIISEVI   223
SaSufD          QYVVLHDDE---------NASFYNHVIIVTEESAEVTYVENYLSNASGEGNQLNIISEVI   221
AthSufD         YLRYFS-GETGDRESKRLPVSNPRVFVLV-EEGGEIGIVEEFVGKDEEGFYWTNPVLEVV   262
EcsufD          LLMHITQGVAGE----EVNTAHYRHHLDL-AEGAEATVIEHFVSLNDARH-FTGARFTIN   206
SfSufD          LLMYITQGVAGE----EVNTAHYRHHLDL-AEGAEATVIEHFVSLNDARH-FTGARFTIN   206
StSufD          LLMHITQGVDGD----ELNTAHYRHHLAL-AEGAEATVIEHYVSLTAAKH-FTGACLTMN   206
                    .  .           .:  .  : :  : ..   :*.::.         .    :

BsSufD          TGDNASVT--YGAVDNLSSGVTTYVNRRGAARGRDSKIEWALGLMNDGDTI-SENTTNLY   280
SaSufD          AGANSNIT--YGSVDYMDKGFTGHIIRRGITE-ADASINWALGLMNEGSQI-IDNTTNLF   277
AthSufD         VQKNAKLKHSYLQKESMASAHIKWTFVRQEAESEYELVEVSTG----GKLGRHNVHVQQL   318
EcsufD          VAANAHLQHIKLAFENPLSHHFAHNDLLLAEDATAFSHSFLLG----GAVLRHNTSTQLN   262
SfSufD          VAANAHLQHIKLAFENPLSHHFAHNDLLLAEDATAFSHSFLLG----GAVLRHNTSTQLN   262
StSufD          VADNAQLRHFKLAFENASSYHFAHNDLLLATDASAFSHSFLLG----AAVLRHHSSSQLN   262
                .  *: :       :   .                   .   *    .     .   :

BsSufD          GDGTYGDTKTVVVGRGEQTENFTTQIIHFGKAS-EGYILKHGVMKDSASSIFNGIGKIEH   339
SaSufD          GDRSTSSLKSVVVGTGEQKINLTSKIVQYGKET-DGYILKHGVMKEHASSVFNGIGYIKH   336
AthSufD         GPDTLTELTTFHMCVNEQTLDLHSKIILDHPRGASRQLHKCIVAHSSGQAVFDGNVRVNR   378
EcsufD          GENSTLRINSLAMPVKNEVCDTRTWLEHNKGFCNSRQLHKTIVS-DKGRAVFNGLINVAQ   321
SfSufD          GENSTLRINSLAMPVKNEVCDTRTWLEHNKGFCNSRQLHKTIVS-DKGRAVFNGLINVAQ   321
StSufD          GENATLRLNSLAMPVKNEVCDTRTWLEHNKGYCNSRQLHKTIVS-DKGRAVFNGLINVAQ   321
                *  :    .:. :   ::  :  : :        .  : *  *  . . ::*:*   : :

BsSufD          GASKANAEQESRVLMLSEKARGDANPILLIDEDDVTAGHAASVGRVDPIQLYYLMSRGIP   399
SaSufD          GGTKSIANQESRVLMLSEHARGDANPILLIDEDDVQAGHAASVGRVDPDQLYYLMSRGIS   396
AthSufD         FAQQTNAGQLTRSLLLKPRATVNIKPNLQIIADDVKCSHGAAISDLEEDQLFYFQARGID   438
EcsufD          HAIKTDGQMTNNNLLMGKLAEVDTKPQLEIYADDVKCSHGATVGRIDDEQIFYLRSRGIN   381
SfSufD          HAIKTDGQMTNNNLLMGKLAEVDTKPQLEIYADDVKCSHGATVGRIDDEQMFYLRSRGIN   381
StSufD          HAIKTDGQMTNNNLLLGKLAEVDTKPQLEIYADDVKCSHGATIGRIDDEQMFYLQSRGIR   381
                 . :: .   .. *::   *  : :* * *  *** ..*.*::. ::  *::*: :***

BsSufD          KEEAERLVIYGFLAPVVNELPIEGVKKQLVSVIERKVK----   437
SaSufD          QREAERLVIHGFLDPVVRELPIEDVKRQLREVIERKVSK---   435
AthSufD         LETARRALISSFGSEVIEKFPNREIRDQARNHVKGLL-----   475
EcsufD          QQDAQQMIIYAFAAELTEALRDEGLKQQVLARIGQRLPGGAR   423
SfSufD          QQDAQQMIIYAFAVELTEALRDEGLKQQVLARIGQRLPGGAR   423
StSufD          QQEARHMILYAFAAELTEAIHDSALKQQVLARIGQRLPGGLV   423
                 . *.: :: .*   : . :    :: *    :   :
```

**Figure S8: Conservation of amino-acids in SufB and SufD proteins.** Multiple-sequence alignments for SufB and SufD constructed using Clustal-Omega software. Amino-acids discussed in this study are underlined in yellow (strict conservation) and in grey (partial conservation). Abbreviations: Bs: *Bacillus subtilis*; Sa: *Staphylococcus aureus*; Ath: *Arabidopsis thaliana*; Ec: *Escherichia coli*; Sf: *Shigella flexneri*; St: *Salmonella typhimurium;* Mm: *Methanosarcina mazei*.

**Figure S9**

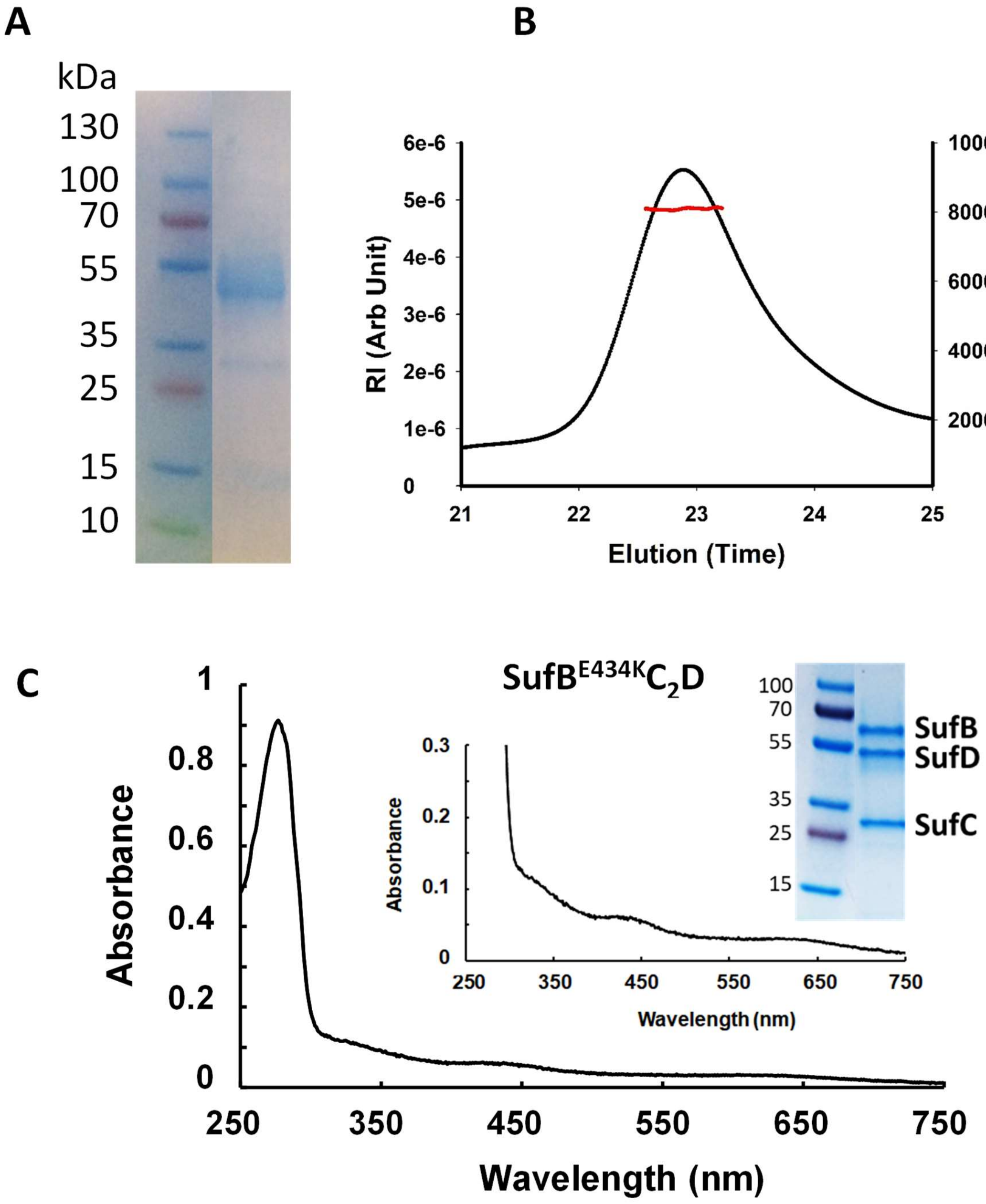


**Figure S9: Biochemical and spectroscopic properties of expressed SufB$^{E434A}$CD and SufB$^{E434K}$CD variants**. SDS-PAGE electrophoresis (A) and SEC-MALS-RI data (B) of the anaerobically purified "SufB$^{E434A}$CD" variant showing that no SufBC$_2$D complex is formed. For SEC-MALS-RI: sample concentration is determined at 2.5 mg.ml$^{-1}$ and gives a MW$_{exp}$ of 83 kDa +/- 0.1%. Differential refractive index RI in Arb Unit (black curve), calculated MW in g/mol (red line). (C) UV-vis. spectrum of SufB$^{E434K}$C$_2$D variant purified anaerobically. Inset: enhancement of the Fe-S absorption bands and SDS-PAGE of the purified SufB$^{E434K}$C$_2$D complex.

#### References

1. Fish, W. W., Rapid colorimetric micromethod for the quantitation of complexed iron in biological samples. *Methods Enzymol.* **1988,** *158*, 357-364.
2. Beinert, H., Semi-micro methods for analysis of labile sulfide and of labile sulfide plus sulfane sulfur in unusually stable iron-sulfur proteins. *Anal. Biochem.* **1983,** *131* (2), 373-378.
3. Proux, O.; Nassif, V.; Prat, A.; Ulrich, O.; Lahera, E.; Biquard, X.; Menthonnex, J. J.; Hazemann, J. L., Feedback system of a liquid-nitrogen-cooled double-crystal monochromator: design and performances. *J. Synchrotron Radiat.* **2006,** *13* (Pt 1), 59-68.
4. Ravel, B.; Newville, M., ATHENA, ARTEMIS, HEPHAESTUS: data analysis for X-ray absorption spectroscopy using IFEFFIT. *J. Synchrotron Radiat.* **2005,** *12* (Pt 4), 537-541.
5. Rehr, J. J.; Kas, J. J.; Prange, M. P.; Sorini, A. P.; Takimoto, Y.; Vila, F., Ab initio theory and calculations of X-ray spectra. *Cr. Phys.* **2009,** *10* (6), 548-559.
6. Newville, M., IFEFFIT: interactive XAFS analysis and FEFF fitting. *J. Synchrotron Radiat.* **2001,** *8* (Pt 2), 322-324.
7. (a) Carboni, M.; Clemancey, M.; Molton, F.; Pecaut, J.; Lebrun, C.; Dubois, L.; Blondin, G.; Latour, J. M., Biologically relevant heterodinuclear iron-manganese complexes. *Inorg. Chem.* **2012,** *51* (19), 10447-10460. (b) Carboni, M.; Molton, F.; Pécaut, J.; Lebrun, C.; Dubois, L.; Blondin, G.; Latour, J. M, *Inorg. Chem.*, **2012**, 51, 12053 (Correction).
8. Charavay, C. Segard, S.; Edon, F.; Clémancey, M.; Blondin, G. *SimuMoss Software*, Univ. Grenoble Alpes, CEA, CNRS, **2012**.
9. Zhang, B.; Bandyopadhyay, S.; Shakamuri, P.; Naik, S. G.; Huynh, B. H.; Couturier, J.; Rouhier, N.; Johnson, M. K., Monothiol glutaredoxins can bind linear [Fe3S4]+ and [Fe4S4]2+ clusters in addition to [Fe2S2]2+ clusters: spectroscopic characterization and functional implications. *J. Am. Chem. Soc.* **2013,** *135* (40), 15153-15164.
10. Krebs, C.; Henshaw, T. F.; Cheek, J.; Huynh, B. H.; Broderick, J. B., Conversion of 3Fe-4S to 4Fe-4S clusters in native pyruvate formate-lyase activating enzyme: Mossbauer characterization and implications for mechanism. *J. Am. Chem. Soc.* **2000,** *122* (50), 12497-12506.
11. Hu, Z.; Jollie, D.; Burgess, B. K.; Stephens, P. J.; Munck, E., Mossbauer and EPR studies of Azotobacter vinelandii ferredoxin I. *Biochemistry* **1994,** *33* (48), 14475-14485.
